# Varigraph: an accurate and widely applicable pangenome graph-based variant genotyper for diploid and polyploid genomes

**DOI:** 10.1101/2025.02.17.638628

**Authors:** Ze-Zhen Du, Jia-Bao He, Pei-Xuan Xiao, Jianbing Hu, Ning Yang, Wen-Biao Jiao

## Abstract

Graph pangenome references can address single-reference bias, thereby enhancing the accuracy of variant genotyping and empowering downstream applications in population genetics and quantitative genetics. However, existing pangenome-based genotyping methods struggle with large or complex pangenome graphs, particularly in polyploid genomes. Here, we introduce Varigraph, an algorithm that leverages the comparison of unique and repetitive *k*-mers between variant sites and short reads for genotyping both small and large variants. Varigraph outperforms current state-of-the-art linear and graph-based genotypers across non-human genomes while maintaining comparable accuracy in human genomes. By employing an efficient data structure, Varigraph achieves higher accuracy in repetitive regions while managing computational costs for large datasets. Notably, Varigraph is the first tool capable of effectively utilizing pangenome graphs for genotyping autopolyploids, enabling precise determination of allele dosage. This work provides a robust and accurate solution for genotyping non-human genomes and will facilitate genomic studies of polyploid crops.

## Introduction

Pangenome graphs, which integrate multiple genome sequences, have emerged as a solution to overcome the bias of single linear reference genomes, thereby enhancing the accuracy of read mapping and variant genotyping^1^. In recent years, several pangenome graph-based tools have been developed for variant genotyping^2–12^, often outperforming traditional single linear reference-based genotypers, especially for structural variations (SVs). These tools have been widely applied in human genomic studies^13^, population diversity analyses, genome-wide association studies (GWAS), and genomic prediction (GS) for plant crops and animals^14–17^.

However, challenges remain for graph-based variant genotyping. First, with advances in long-read sequencing, more genomes are now *de novo* assembled, leading to larger and more complex pangenome graphs. This requires rapid and memory-efficient algorithms and data structures. More graphed genomes typically introduce more highly similar allelic variants, increasing the number of multi-branch nodes in the graph. These complex regions challenge the accuracy of variant genotyping.

Second, most graph-based genotyping tools are tailored for human genomes and struggle with plant genetic variants^18^. This is due to the intrinsic features of plant genomes, such as excessive repetitive sequences, variable genome size, and high heterozygosity. For example, while the rice genome (373 Mb) is much smaller than the human genome (3,055 Mb), it has a higher ratio of repetitive *k*-mers^19^. Not to mention other plants such as wheat (*Triticum aestivum*), which possesses an exceptionally large genome size of approximately 14.5 Gb and an even higher proportion of repetitive sequences. This can significantly decrease genotyping performance, especially for tools that rely solely on unique *k*-mers^8, 12^. Additionally, there are no available graph-based genotypers for polyploid genomes. Many important crops, such as wheat, potato, rapeseed, and sugarcane, are polyploid, necessitating sophisticated genotyping models and algorithms for these genomes.

In this study, we developed Varigraph, a pangenome graph-based genotyping tool for small (SNPs and indels) and large (SVs) variants from both diploid and polyploid genomes. Varigraph directly compares *k*-mers at variant loci and short reads without read mapping, accelerating the genotyping process. In particular, it also considers non-unique *k*-mers for complex regions to improve genotyping efficiency. By implementing Counting Bloom Filter and bitmap data structures, Varigraph can handle super pangenome graphs with more than 200 genomes. Importantly, it supports genotyping in polyploid genomes using a modified version of the Hidden Markov Model. We evaluated and compared the performance of Varigraph with other state-of-the-art genotyping tools on the human genome and a series of representative plant genomes. These comparisons demonstrated Varigraph’s superiority in nearly all scenarios, including challenging cases such as genotyping variants in repetitive regions, highly repetitive genomes, large pangenome graphs, and polyploid genomes.

## Results

### Overview of Varigraph algorithm

The workflow of Varigraph algorithms mainly includes five steps **(Fig. 1)**. Based on a reference genome and variants of alternative genomes, Varigraph starts by constructing a directed acyclic variation graph (step 1). In this graph, each node represents sequences, each edge between nodes indicates the connection of sequences, and the allelic variants are shown as bubbles. Additionally, Varigraph uses a Counting Bloom Filter to store all *k*-mers in the reference genome (step 2). This data structure enables storing all *k*-mers in the genome with a memory footprint approximately tenfold that of the genome itself, while offering constant-time complexity for querying the frequency of any given *k*-mer. In contrast, another *k*-mer based graph genotyper, BayesTyper, uses a Bloom Filter without counting functionality, which limits its effectiveness in handling k-mer frequency information.

**Fig. 1.**
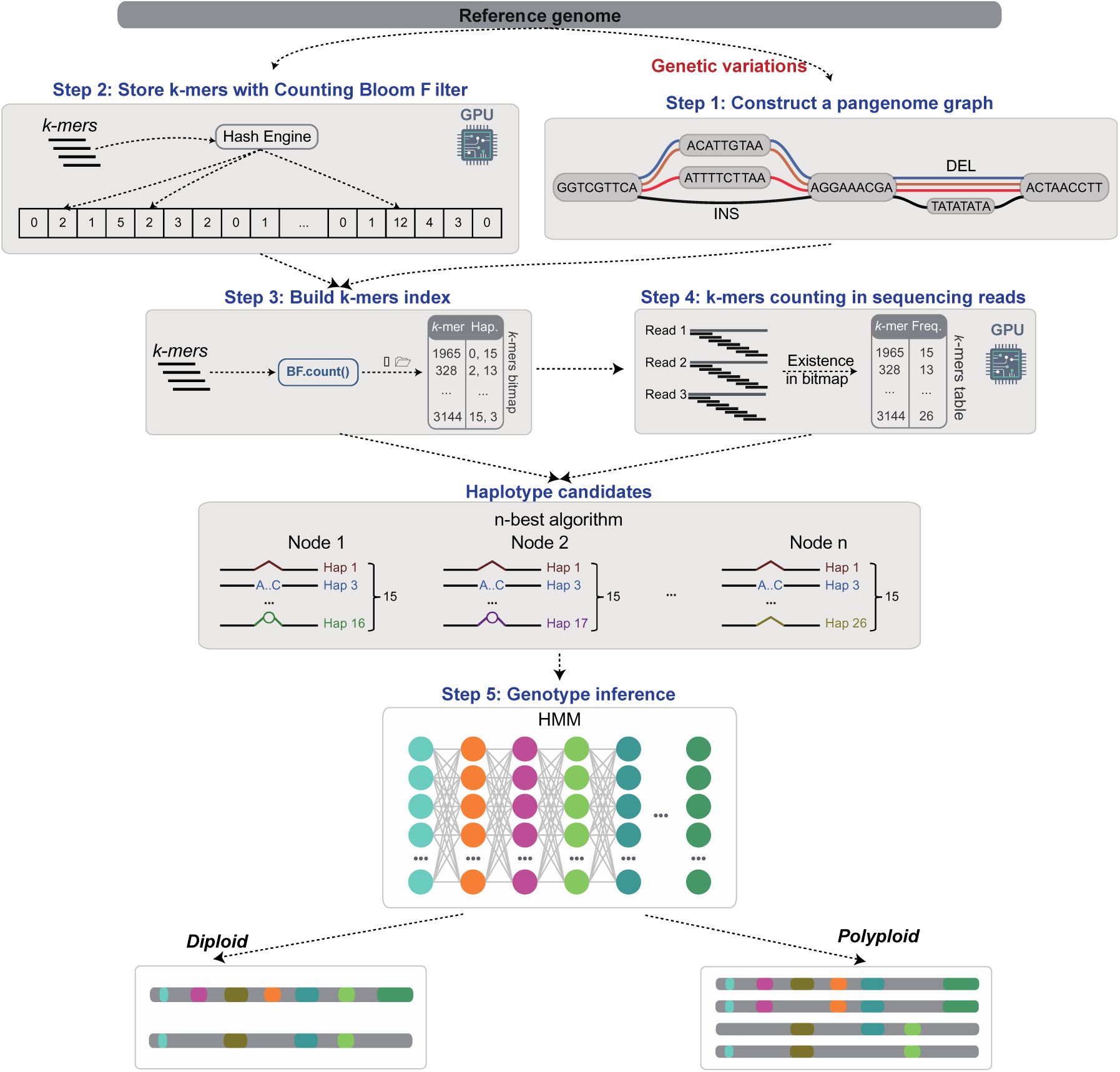
Overview of the Varigraph algorithm. The whole workflow includes five steps. Step 1: a pangenome graph is constructed using the reference and variants from alternative genomes. Step 2: Varigraph constructs a Counting Bloom Filter based on the reference genome. It also provides a GPU-accelerated mode for this construction. Step 3: Varigraph indexes the pangenome graph by *k*-mers, storing their information in a hash table with bitmap structure. Step 4: Varigraph counts the frequence of *k*-mers present in the genome graph within the reads. It also supports a GPU-accelerated mode. Step 5: The most likely haplotypes (default 15) are selected based on *k*-mers frequence and used for genotype inference. By modeling the *k*-mers frequence with a Hidden Markov Model (HMM), Varigraph infers variant genotypes. Two different models are used for genotyping diploid and autopolyploid genomes, respectively.

Varigraph then builds a *k*-mer index for each node in the genome graph (step 3). Other graph based genotypers, such as BayesTyper and PanGenie, that only load unique *k*-mers from the genome graph for variant genotyping. However, unique *k*-mers might be not sufficient when the genome enriches repetitive sequences, or the graph included many variants from repetitive regions. To mitigate this issue, Varigraph combines non-unique *k*-mers to enhance the accuracy of genotyping. Furthermore, to minimize memory usage, Varigraph stores *k*-mers as 64-bit integers, allowing for a maximum *k*-mer length of 28. In addition, Varigraph employs a bitmap data structure to store haplotype information for each *k*-mer, which reduces the memory consumption to 1/16th of the original size (Methods). More importantly, it supports a pangenome graph that can accommodate up to 65,535 haplotypes, equivalent to 32,767 alternative diploid genomes. In contrast, PanGenie, which uses 8-bit integers to store haplotype information, is limited to supporting only 255 haplotypes.

Then, Varigraph scans *k*-mers of sequencing reads of each individual to be genotyped (step 4). It subsequently locates the corresponding *k*-mers in the genome graph index and updates their depth values, and the depth information for each *k*-mer which is stored as an 8-bit integer to reduce the memory usage. Besides, Varigraph introduces a GPU-accelerated mode for the construction of Counting Bloom Filter with speedups 3.4 to 47 times, while also for sequencing data traversal by speedup of approximately 2.5 times for five genomes **(Supplementary Table 1)**.

Subsequently, Varigraph calculates the probability of occurrence of each haplotype in the population based on the *k*-mers depths and selects the top 15 haplotypes (by default) for genotyping. For diploid, Varigraph computes the prior probability of each haplotype pair combination based on the *k*-mer frequencies contained in the haplotypes and their depth in the sequencing data. It then uses a Hidden Markov Model (HMM) to calculate the probability of each haplotype pair combination at each variant site. Finally, Varigraph outputs the genotype corresponding to the haplotype combination with the highest posterior probability (step 5). For the genotyping of autotetraploid genomes, Varigraph implements a similar approach by calculating the posterior probability of any four haplotype combinations and outputs the genotyping information of the optimal combination (Methods).

### Constructing datasets for comparisons of genotyping performance

To evaluate the performance of Varigraph against state-of-the-art pangenome graph-based genotypers, we performed variant genotyping tests using both simulated and real datasets constructed based on long-read assemblies, third-generation long sequencing reads, and second-generation short sequencing reads (Methods). Our evaluations encompassed the human genome and a range of plant diploid and polyploid genomes **(Supplementary Table 2)**. Pangenome graphs were constructed by incorporating one reference genome and variants derived from alternative genomes **(Supplementary Tables 3-4)**. For *A. thaliana* and rice, genomes with known ground-truth variants were used to simulate short reads for genotyping. These ground-truth variants were simulated based on variants called from real genomes **(Supplementary Table 5; Methods)**. To identify high-quality variants from real genomes for pangenome graph construction and performance benchmarking, we developed a pipeline by integrating assembly comparisons and alignments of long and short reads **(Supplementary Fig. 1)**. For genotyping in other species, we used real short reads from genomes where their long-read assemblies and original long-reads were also available for identification of benchmarking variants **(Supplementary Table 6)**.

### Comparisons to existing methods on human genomes

We firstly compared Varigraph with other graph-based variant genotypers using a human pangenome graph constructed based on the Human Pangenome Reference Consortium (HPRC) dataset. This dataset comprises 45 haplotype-assembled human genomes, encompassing 19,104,858 SNPs, 7,836,674 indels (< 50 bp), 93,746 deletions (>= 50 bp), and 333,597 insertions (>= 50 bp) **(Supplementary Table 3)**. We run genotyping for each tool using short reads from three human samples. Tools like vg map, vg giraffe were not tested due to computational limitation under such a large number of variants. Similarly, GraphTyper2 was only evaluated for genotyping insertions and deletions.

Overall, Varigraph displayed performance very close to that of the top-performing genotyper, PanGenie **(Fig. 2, and Supplementary Table 7)**. For variants in nonrepetitive regions, Varigraph had the same or slightly lower F-scores compared to PanGenie. For example, both Varigraph and PanGenie presented F-scores of 0.96 for SNP genotyping. For variants in repetitive regions, their F-scores was even close, with an average difference only 0.03. Notably, Varigraph achieved a higher F-score (0.74) than PanGenie (0.72) for genotyping deletions in repetitive regions **(Fig. 2j)**. Additionally, Varigraph showed F-scores comparable to PanGenie for both bi-allelic and multi-allelic variants **(Supplementary Fig. 2)**. Additionally, we evaluated the performance of Varigraph using short-read datasets with varying sequencing depths. With 10X coverage, Varigraph attained F-scores of 0.72-0.90 (SNPs: 0.90, small indel: 0.88, midsize indels: 0.81, insertions: 0.80, deletions: 0.72). It reached nearly optimal genotyping performance when 20X coverage was provided **(Supplementary Fig. 3)**. Similar sequencing depth requirements were observed for PanGenie. Furthermore, both Varigraph and PanGenie showed sensitivity to breakpoint errors in structural variants (SVs) **(Supplementary Fig. 4)**.

**Fig. 2.**
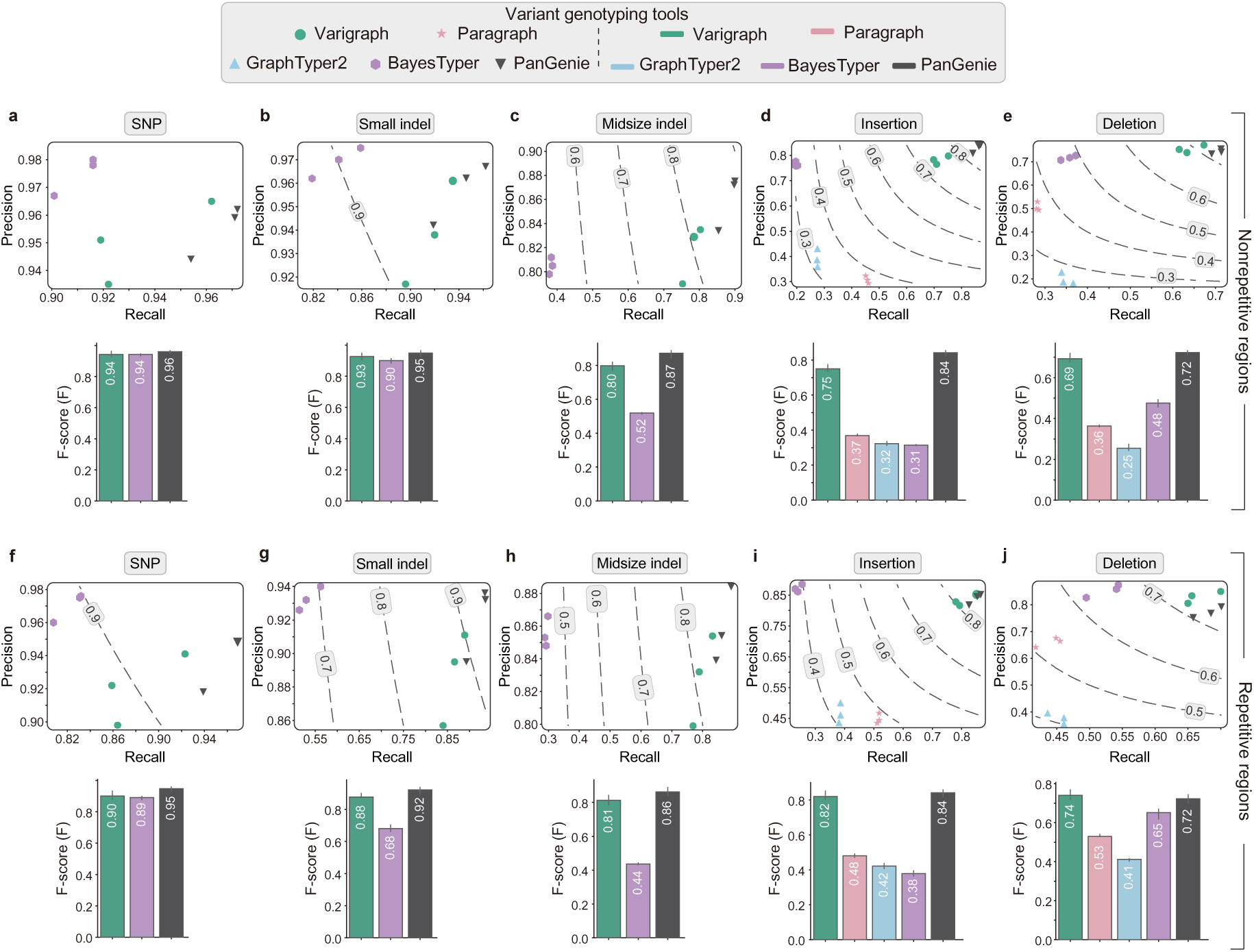
Performance comparison of Varigraph and other pangenome graph-based genotypers in variant genotyping for human genomes. Genotyping performance for SNPs, small indels (1-19 bp), midsize indels (20-49 bp), insertions (>=50 bp), and deletions (>=50 bp) in nonrepetitive **(a-e)** and repetitive regions **(f-j)** of human genomes. The genome graph was constructed with one reference genome and 45 alternative genomes from the Human Pangenome Reference Consortium (HPRC) dataset. Paired-end (2 × 149 bp) short reads with depths of 35.9× for HG00438, 35.9× for HG01928and 37.9× for NA20129 were used for genotyping (as shown with three symbols in the figure for each genotyper). The dashed lines correspond to recall and precision under different F-scores. Numbers on the dashed lines indicate F-scores. The histograms represent the average F-scores for genotyping of three genomes.

### Performance on representative diploid plant genomes

We initially compared the performance of Varigraph and other genotypers on an *A. thaliana* pangenome graph. This pangenome graph was constructed based on the reference genome Col-0 accession and variants from high-quality genome assemblies of 65 accessions **(Supplementary Table 3)**. Variants include 4,139,626 SNPs, 662,839 indels, and 115,223 SVs. One genome was simulated by random selection of these variants and 30X paired-end (2 x150 bp) short reads was produced from this genome and used for variant genotyping.

For nonrepetitive regions, Varigraph consistently achieved the highest F-scores for all variant types (SNPs: 0.99, small indels: 0.94, midsize indel: 0.97, deletions: 0.95, insertions: 0.98) **(Fig. 3a, Supplementary Fig. 5 and Supplementary Table 8)**. In repetitive regions, although Varigraph’s attained the highest F-scores for SNPs (0.91), midsize indels (0.84), insertions (0.95), and deletions (0.91) **(Fig. 3b, Supplementary Fig. 5 and Supplementary Table 8)**. Compared to genotyping non-repetitive regions, Varigraph showed the smallest decrease in F-scores for genotyping in repetitive regions, whereas other tools suffered more drops. Notably, the k-mer based genotyping tools BayesTyper and PanGenie exhibited the most substantial performance reductions in repetitive regions, likely due to their algorithms’ reliance on unique k-mers.

**Fig. 3.**
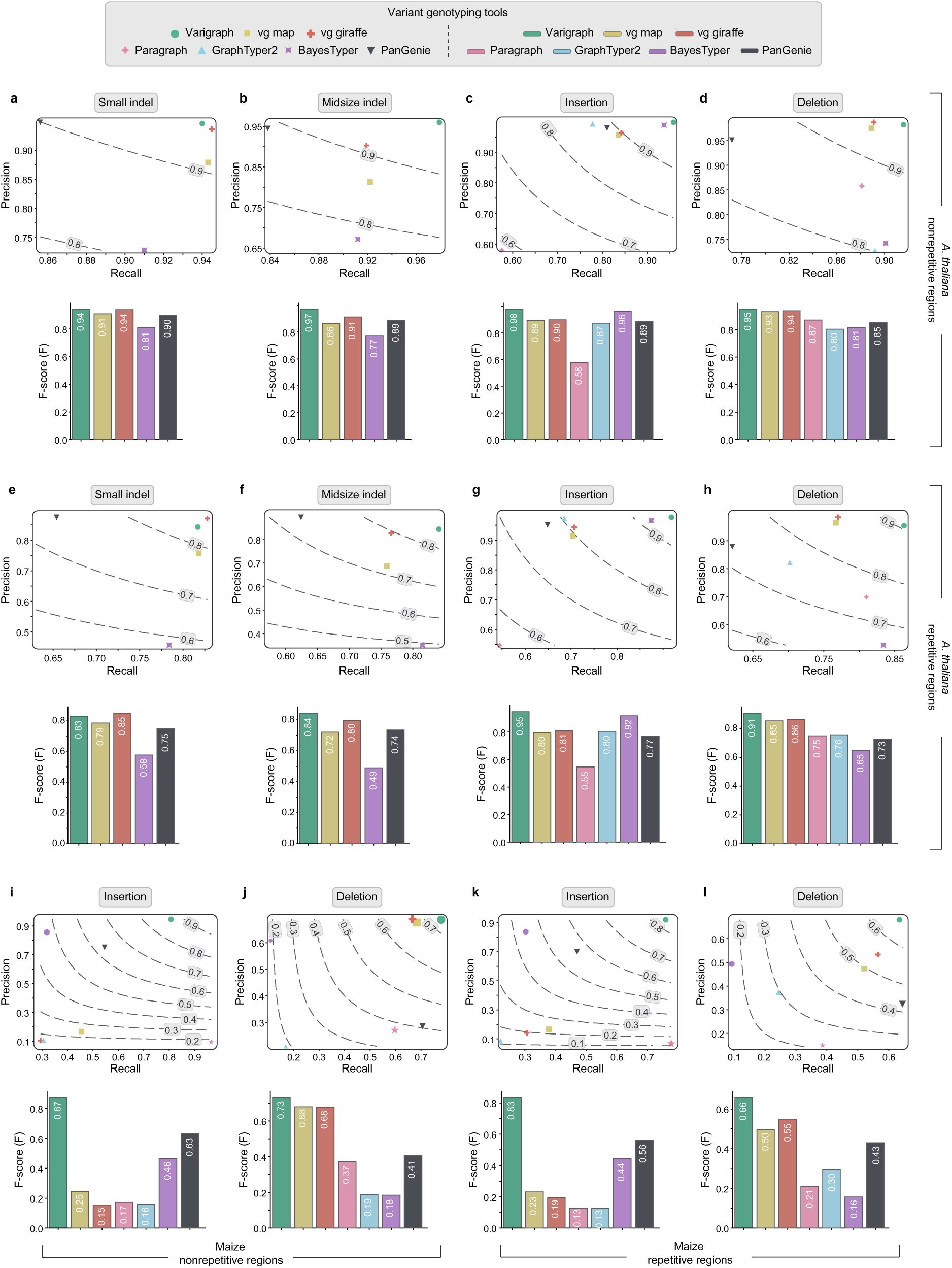
Performance comparison of Varigraph and other pangenome graph-based genotypers in variant genotyping for diploid plant genomes. **(a-h)** Genotyping performance for small indels (1-19 bp), midsize indels (20-49 bp), insertions (>=50 bp), and deletions (>=50 bp) in nonrepetitive **(a-d)** and repetitive regions **(e-h)** of *A. thaliana* genomes. **(i-l)** Genotyping performance for insertion and deletions in nonrepetitive **(i, j)** and repetitive regions **(k, l)** of maize genomes. The dashed lines correspond to recall and precision under different F-scores. Numbers on the dashed lines indicate F-scores.

To investigate the impact of heterozygous genomes on graph-based genotypers, we performed testes on a dataset consisting of 66 synthetic heterozygous *A. thaliana* genomes including 4,241,614 SNPs, 668,408 indels, and 118,466 SVs **(Supplementary Table 3)**. Again, Varigraph achieved the best performance for genotyping in nonrepetitive regions (F-scores: 0.98, 0.92, 0.95, 0.91, 0.91 for SNPs, small indels, midsize indels, deletions, and insertions, respectively, **Supplementary Fig. 6 and Supplementary Table 9**). This performance was close to that observed in homozygous *A. thaliana* genomes. In contrast, alignment-based graph genotypers such as vg map, vg giraffe, and GraphTyper2 showed considerably decreased performance in genotyping SVs on this heterozygous genome dataset. In repetitive regions, the F-scores for all tools decreased by approximately 0.1. Overall, Varigraph outperformed all other tools for this dataset, except for small indels in repetitive regions where Varigraph had a F-score very closed to the best, vg giraffe (0.79 vs 0.81) **(Supplementary Fig. 6 and Supplementary Table 9)**.

We further performed variant genotyping on real heterozygous diploid pummelo (*Citrus maxima*) genomes **(Supplementary Fig. 7 and Supplementary Table 10)**. We constructed a pangenome graph with 8 genomes including 5,186,044 SNPs, 1,412,716 indels, and 96,303 SVs. In non-repetitive regions, Varigraph had the highest F-score for deletions (0.71), while its F-scores in genotyping other types of variants was on par with the top performers, exhibiting only a marginal average difference of 0.02. In repetitive regions, Varigraph excelled in genotyping indels, insertions, and deletions. Notably, the genotyping performance of other tools in repetitive regions was significantly affected, with an average F-score drop of 0.13 compared to non-repetitive regions. In contrast, Varigraph demonstrated the most stable performance, with only an average decrease of 0.08.

In addition, we conducted tests on the highly repetitive maize (*Zea mays*) genome, which consists of 89% repeats. We constructed a maize pangenome graph incorporating 25 genomes with 1,161,109 insertions and 580,315 deletions **(Supplementary Table 2)**. Due to the computational limitation, SNPs and indels were not included in the graphs. Varigraph showed the highest F-score for insertions (0.87) and deletions (0.73) at non-repetitive regions and considerably outperformed other genotyper for variants at repetitive regions **(Fig. 3i-l, and Supplementary Table 11)**. Other graph-based genotypers only performed well in only one type of variant or only in non-repetitive regions. For example, vg map had a F-score of 0.68 for genotyping nonrepetitive deletions but only 0.25 for nonrepetitive insertions. Overall, Varigraph consistently showed the best performance in genotyping these plant genomes compared to other genotypers.

### Performance on large-scale pangenome graphs

Increasing the number of graphed genomes expands the search space for read alignment or *k*-mer comparisons, leading to higher computational cost and often reducing performance of graph-based genotyping tools^18^. To further evaluate Varigraph’s performance on large pangenome graphs, we used a dataset comprising 252 diploid homozygous rice genomes, which included 15,177,337 SNPs, 1,828,713 indels, and 803,284 SVs **(Supplementary Table 3)**. Due to algorithmic limitations, PanGenie was tested on only 127 rice genomes. Other genotypers, such as vg map, vg giraffe, Paragraph, and GraphTyper2, were tested only for SV genotyping due to running failures or computational limitations.

In nonrepetitive regions, Varigraph presented the highest F-scores for all variant types (0.99, 0.95, 0.99, 0.98, 0.94 for SNPs, small indels, midsize indels, insertions, deletions, respectively) **(Fig. 4a-d, Supplementary Fig. 8, and Supplementary Table 12)**. In repetitive regions, Varigraph showed only a minor drop in F-scores and remained the best performance for all types of variants **(Fig. 4e-h, Supplementary 9, and Supplementary Table 12)**. For example, Varigraph maintained high F-scores even for large variants (insertions: 0.92, deletions: 0.90). However, all other genotyping tools had relatively lower F-scores, indicating the advantage of Varigraph in large pangenome graphs and repetitive regions.

**Fig. 4.**
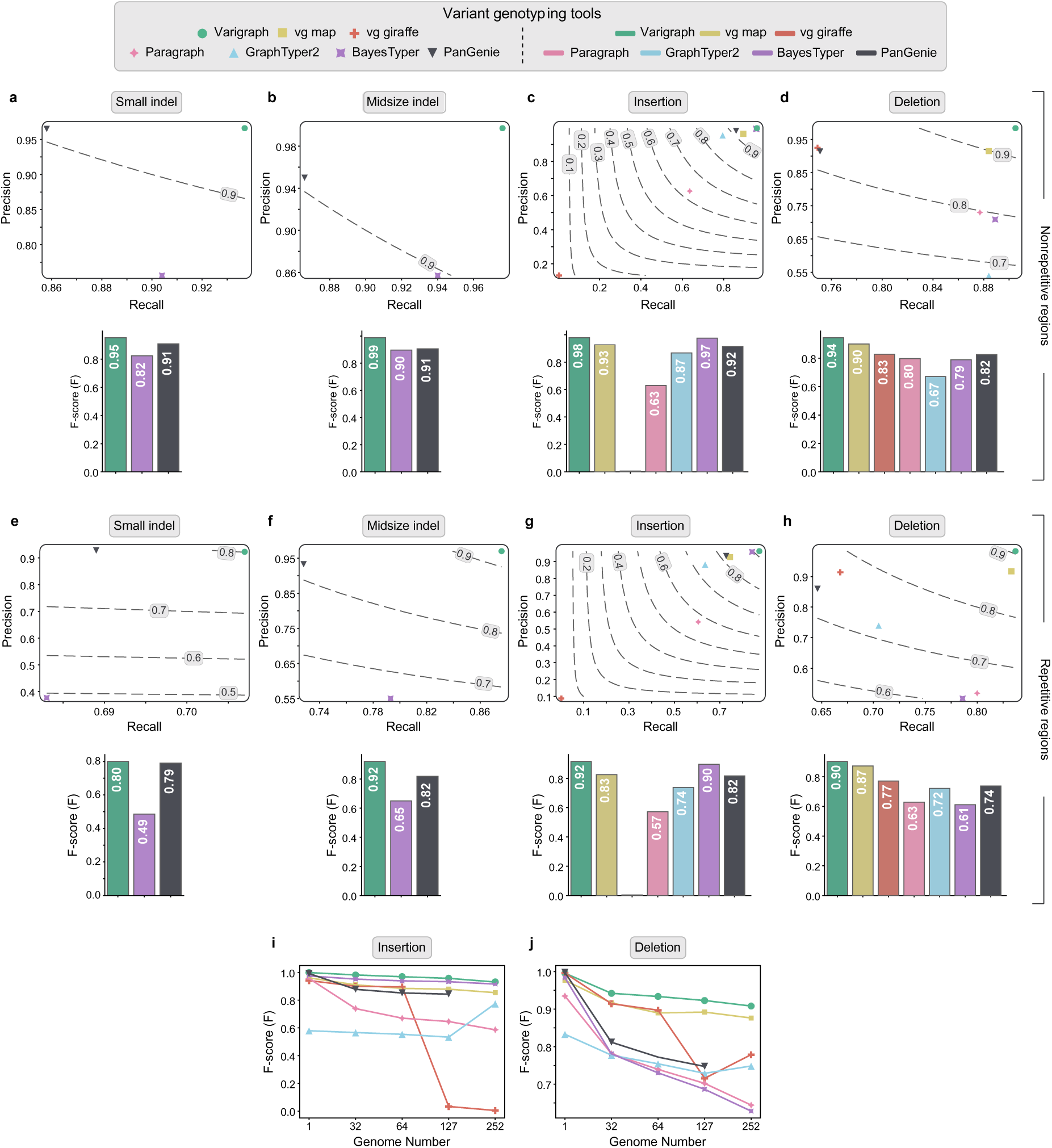
Performance comparison of Varigraph and other pangenome graph-based genotypers in variant genotyping for a large pangenome graph with 252 rice genomes. **(a-h)** Genotyping performance for small indels (1-19 bp), midsize indels (20-49 bp), insertions (>=50 bp), and deletions (>=50 bp) in nonrepetitive **(a-d)** and repetitive regions **(e-h)**. The dashed lines correspond to recall and precision under different F-scores. Numbers on the dashed lines indicate F-scores. **(i, j)** Genotyping F-scores for insertions **(i)** and deletions **(j)** under different number of graphed genomes. Note: PanGenie was tested on 127 rice genomes due to the algorithm limitation while other genotypers, vg map, vg giraffe, Paragraph, and GraphTyper2 were tested only for SVs genotyping due to running failure or computational limitation.

To further investigate the impact of the number of genomes on variant genotyping performance, we repeated the tests on a series of pangenome graphs integrated with 1, 32, 64, 127, and 252 alternative genomes, respectively. As the number of genomes increased, Varigraph demonstrated the highest stability in performance. Specifically, under the integration of 252 genomes, Varigraph’s F-scores for SNPs, indels, deletions, and insertions were 94%, 85%, 91%, and 93%, respectively, compared to those obtained with two genomes **(Fig. 4k, l, Supplementary Fig. 9, and Supplementary Table 13)**. In contrast, other tools suffered significant decreases in genotyping performance as more genomes were graphed. For example, vg giraffe was unable to genotype insertions when all 252 genomes were included.

### Performance on allopolyploid genomes

Next, we evaluated the genotyping performance of Varigraph in allopolyploid genomes using a pangenome graph of seven allotetraploid *Brassica napus* (2n=4x=38) genomes, which included 11,662,049 SNPs, 3,609,178 indels, and 284,648 SVs **(Supplementary Table 2)**. Varigraph employs a diploid genotyping model for allopolyploids which have considerable sequence divergence between subgenomes. Compared to other graph-based genotypers, Varigraph always presented the highest F-scores for all types of variants from nonrepetitive regions (SNPs: 0.92, small indels: 0.88, midsize indels: 0.90, insertions: 0.65, deletions: 0.80) and repetitive regions (SNPs: 0.80, small indels: 0.76, midsize indels: 0.82, insertions: 0.56, deletions: 0.78) **(Supplementary Fig. 10 and Supplementary Table 14)**. Besides, Varigraph showed the smallest difference in genotyping performance between nonrepetitive and repetitive regions.

Additionally, we tested Varigraph on the huge allohexaploid genome of bread wheat (14.5 Gb). A pangenome graph was build based on 346,833 indels and 47,918 SVs from 10 wheat genomes **(Supplementary Table 3)**. Varigraph took 4.4 hours for pangenome graph construction with a memory peak of 156 Gb, and 25 hours for variant genotyping with a memory peak of 16 Gb. It presented F-score over than 0.7 for all types of variants **(Supplementary Fig. 11)**. Other tools were not tested due to computational resource limitations.

### Performance on autopolyploid genomes

Varigraph conducted genotyping in autopolyploid genomes using an autopolyploid model. For example, for a biallelic variant (e.g.: A/a) in the autotetraploid genomes, Varigraph calculates the genotyping probabilities for all five possible genotypes (AAAA, AAAa, AAaa, Aaaa, aaaa). Since no other graph-based genotyping tools are available for autopolyploids, we compared Varigraph’s genotyping performance on SNPs and indels with three linear reference-based polyploid genotypers: GATK, FreeBayes, and Octopus.

We first evaluated the genotyping performance based on a pangenome graph including 33 synthetic autotetraploid *A. thaliana* genomes with 4,241,614 SNPs, 668,408 indels, and 118,466 SVs **(Supplementary Table 3)**. For variants in nonrepetitive regions, Varigraph displayed the highest F-scores for all types of variants (SNPs: 0.98, small indels: 0.98, midsize indels: 0.93, insertions: 0.94, deletions: 0.96) **(Supplementary Fig. 12 and Supplementary Table 15)**. For genotyping in repetitive regions, Varigraph still achieved the highest F-scores, only except for SNP genotyping (Varigraph: 0.91, Octopus: 0.93).

Furthermore, we conducted tests of genotyping variants for real autopolyploid genomes. We constructed a pangenome graph using ten published potato genomes which included 58,925,315 SNPs, 11,023,430 indels, 389,575 deletions, and 416,897 insertions **(Supplementary Table 3)**. We run the variant genotyping with Illumina short reads from two autotetraploid potato genomes, Otava^20^ and C88^21^ whose haplotype-resolved long-read assemblies had been published previously. Due to the computational limitation, FreeBayes and Octopus failed to complete the genotyping.

For SNPs and indels from nonrepetitive or repetitive regions, Varigraph consistently showed much higher F-scores compared to GATK **(Fig. 5a-b and Supplementary Table 16)**. For example, Varigraph achieved F-scores of 0.85 (SNPs), 0.78 (small indel), and 0.79 (midsize indel), while GATK had F-scores of 0.75, 0.56, and 0.38, respectively. Additionally, Varigraph demonstrated more stable performance in genotyping variants from repetitive regions, compared to GATK. Furthermore, Varigraph completed the genotyping for one sample in less than two hours, while GATK took more than three days, even when running in parallel at the chromosome level. Besides, Varigraph maintained F-scores ranging from 0.73 to 0.81 for genotyping insertions (repetitive: 0.79, non-repetitive: 0.81) and deletions (repetitive: 0.73, non-repetitive: 0.75). It also displayed stable F-scores for insertions and deletions larger than 1 kb **(Supplementary Fig. 13)**.

**Fig. 5.**
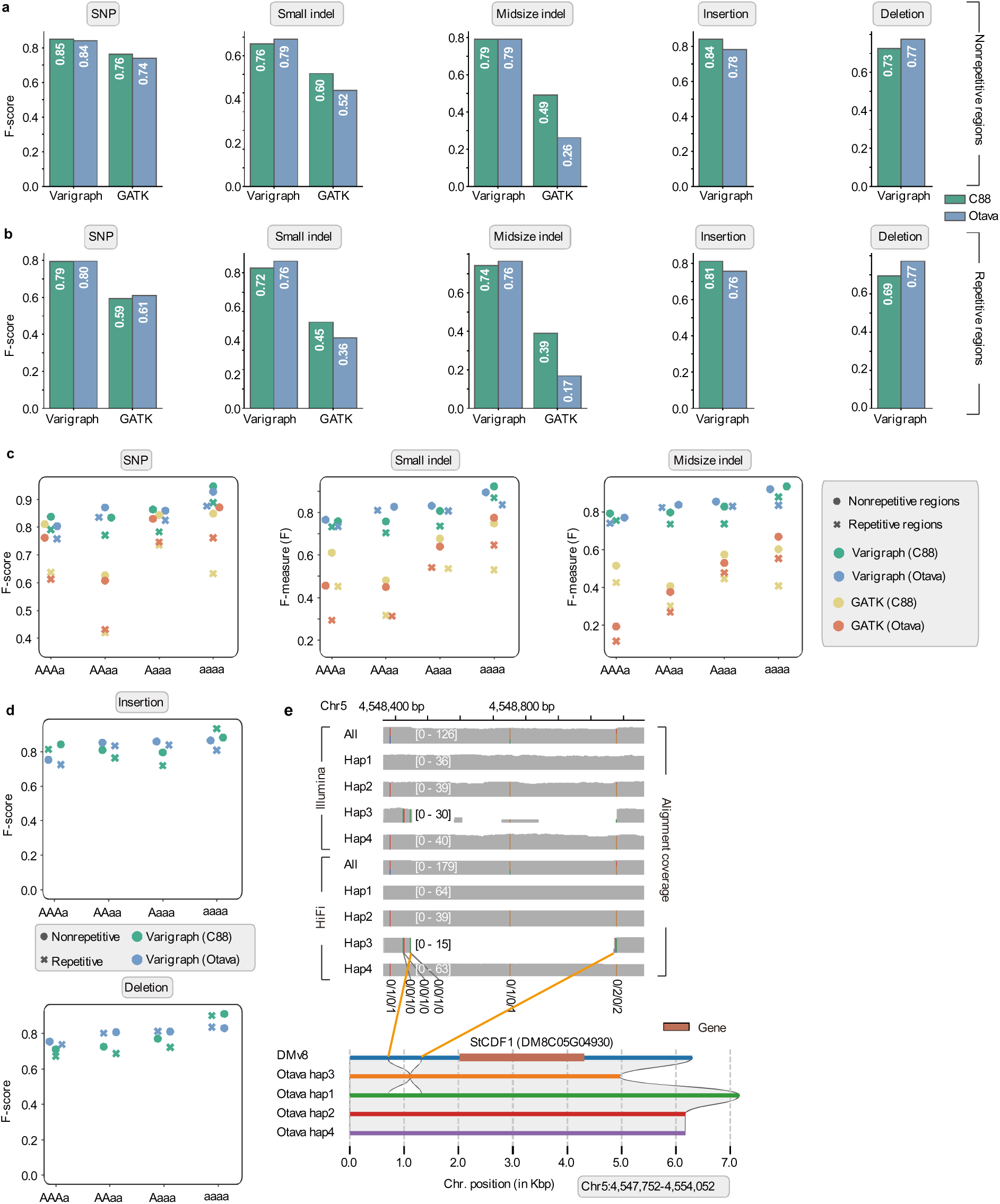
Genotyping performance for variants in autopolyploid potato genomes. **(a-b)** F-score for genotyping SNPs, small indels (1-19 bp), midsize indels (20-49 bp), insertions (>=50 bp) and deletions (>= 50 bp) from nonrepetitive **(a)** and repetitive **(b)** genomic regions. **(c-d)** F-scores for genotyping different genotypes of bi-allelic variants from repetitive regions and nonrepetitive regions of potato genomes (two cultivars: C88 and Otava). Both Varigraph and GATK were run for SNP, small indels (1-19 bp), and midsize indels (20-49 bp), while only Varigraph supports genotyping insertion (>=50 bp) and deletions (>= 50 bp). **(e)** An example at locus of gene *StCDF1* showing genotyping for variants with different dosage. The top panel shows the alignment coverage of Illumina short reads and PacBio HiFi long reads at each haplotype (Hap1, Hap2, Hap3, Hap4) and the total genomic region. Colored bars indicate small variants. The blank region of coverage indicates deletions. The bottom panel shows the sequence alignments between haplotypes of Otava and potato reference genome DMv8 and indicates a heterozygous deletion at the upstream of gene *StCDF1*, which was correctly genotyped by Varigraph.

Remarkably, Varigraph presented more stable performance on different genotypes of bi-allelic variants for both small and large variants **(Fig. 5c-d and Supplementary Table 17)**, suggesting that it can efficiently distinguish different allele dosages. For example, six heterozygous SNPs and one heterozygous deletion were within the gene *StCDF.1*, which regulates plant maturity and initiation of tuber development^22^. Varigraph accurately determined the genotype for all variants at this region, supporting by both haplotype assemblies and alignments of phased reads **(Fig. 5e).** However, GATK was unable to genotype the deletion and failed to call correct genotypes of three SNPs, perhaps due to the biased coverage profiles when only a single linear reference was used.

Compared to bi-allelic variant genotyping, our tests on Varigraph revealed only a 0.07 decrease on average F-scores for multi-allelic SNP and indel genotyping, while GATK showed a 0.19 decrease in such F-scores **(Supplementary Fig. 14)**. Remarkably, Varigraph presented F-scores of 0.74-0.82 for multi-allelic insertions and deletions. In addition, Varigraph demonstrated better genotyping performance when the sequencing coverage per haplotype was between 10 X and 20 X **(Supplementary Fig. 15)**. More sequencing data were not able to improve the genotyping of Varigraph, a finding that is consistent with previous observations for GATK^23^.

### Runtime and memory cost

Additionally, Varigraph demonstrated high efficiency in terms of runtime and memory peak usage **(Supplementary Fig. 16)**. In tests on the datasets of maize and pummelo, Varigraph was 1-2 orders of magnitude faster than the read-mapping-based genotyper, vg map. Furthermore, it run faster than BayesTyper, a *k*-mer-based pangenome graph genotyper, despite being marginally slower than PanGenie. Notably, PanGenie achieved its speed advantage by loading only unique *k*-mers during genotyping. Moreover, for genotyping in autotetraploid potato genomes, Varigraph could complete the genotyping for one sample in less than 4 hours using 10 threads. Similarly, Varigraph showed comparable memory usage to other graph-based genotypers.

## Discussion

In this study, we developed Varigraph, a pangenome graph-based variant genotyper capable of handling both small and large variants in diploid and polyploid genomes. By employing *k*-mer comparisons, Varigraph bypasses the time-consuming process of read alignments. Unlike other *k*-mer-based genotypers, Varigraph loads both unique and non-unique *k*-mers when necessary. Consequently, Varigraph outperforms other graph-based genotypers in genomes enriched with repetitive *k*-mers, especially for variants in repetitive regions and multi-allelic sites. To mitigate potential increases in memory usage due to storing non-unique *k*-mers, it additionally utilizes data structures of Counting Bloom Filter and bitmap.

Varigraph maintained comparable performance in human genome with the best genotyper, PanGenie, which leverages longer linkage disequilibrium information and recombination rates to strengthen genotyping efficiency. However, such data may not always be available or sufficiently accurate for all species. In cases where unique *k*-mers are unavailable, PanGenie estimates genotyping probabilities based on previous loci and recombination rates. This can propagate errors in highly repetitive genomes, leading to decreased performance. In contrast, Varigraph demonstrates superior performance in non-human genomes, particularly in plant genomes with high repetitiveness.

As more *de novo* assemblies become available for pangenome studies, efficient methods for pangenome graph construction and variant genotyping will be essential. Varigraph excels in large pangenome graphs, making it a promising choice for future super-pangenome analyses. It supports giant pangenome graphs, such as those including 252 rice genomes or 10 large wheat (14 Gb) genomes, with memory usage under 356 Gb. Other tools often struggle with computational limitations or support only limited graph sizes. For instance, the current version of PanGenie is restricted to a maximum of 255 haplotypes.

To our knowledge, Varigraph is the first tool to perform graph-based variant genotyping for autopolyploid genomes. It can accurately distinguish different heterozygous genotypes in autopolyploids, which is crucial for allele dosage-based analyses such as genome-wide association studies and genome prediction^24, 25^. Additionally, it also performs well in allopolyploid genomes, even in large and highly repetitive hexaploidy wheat genomes. This makes it a promising tool for variant-based population genetic and quantitative genetic studies in many other important crops such as rapeseed, cotton, and potato.

While Varigraph consistently shows superior performance, further improvements are needed, particularly for complex structural variants (SVs) such as inversions, duplications, and composite SVs. Additionally, new algorithms and models are required to handle more complex polyploids, such as segmental polyploids that contain both allo- and auto-subgenomes. Although we only have tested Varigraph in human and plants, it theoretically should be applicable for animal and other genomes.

In summary, we have developed an accurate pangenome graph-based variant genotyping tool, suitable for genomes of varying complexities, especially autopolyploid genomes and large pangenome graphs. We believe our method will be widely applied for genotyping and downstream studies, such as association studies and phenotype prediction.

## Methods

### Collection of genome assemblies and sequencing reads

The evaluation encompassed human genomes and plant species with varying genome complexities in term of genome size, repeat sequence content, and ploidy. These plant species included the simple homozygous diploid plant genomes of *Arabidopsis thaliana* and rice (*Oryza sativa*), the high-repeat genome of maize, the heterozygous diploid genome of pummelo (*Citrus maxima*), the allotetraploid genome of rapeseed (*Brassica napus*), the autotetraploid potato (*Solanum tuberosum*), and the large allohexaploid genome of bread wheat (*Triticum aestivum*) **(Supplementary Table 2)**. The dataset of each species includes long-read *de novo* assemblies of multiple genomes, the long sequencing reads that were used for each assembly and short reads from the same samples. Details of each dataset were described as below.

#### A. thaliana

Long-read *de novo* genome assemblies of 65 accessions were published in a previous study^26^. The reference genome was the Col-0 TAIR10 version^27^.

#### Rice

Long-read *de novo* genome assemblies of 251 samples were from a previous study^28^.

The reference genome was the line Nipponbare (IRGSP-1.0)^29^.

#### Maize

Long-read *de novo* genome assemblies of 26 lines were published in a previous study^30^. The reference genome was from the line B73 (version 5). The second- and third-generation sequencing reads of these lines were downloaded from the ENA BioProject database, under accession numbers PRJEB31061 and PRJEB32225.

#### Pummelo

The PacBio HiFi sequencing reads of seven *Citrus maxima* cultivars (C5, C9, C31, C34, C35, C39, and GBY) were generated by our group and have been deposited into the National Genomics Data Center (NGDC) database (BioProject ID PRJCA035310) and China National GeneBank DataBase (BioProject ID CNP0006728). Additionally, the short reads from the GBY sample were downloaded from the NCBI database (Accession Number: SRR3823447) and used for variant genotyping.

#### Rapeseed

Long-read *de novo* genome assemblies of sever cultivars were published in a previous study^31^. The reference genome was from the cultivar ZS11. The PacBio SMRT and Illumina sequencing reads of these cultivars were downloaded from the NCBI database under accession number PRJNA546246.

#### Potato

This dataset included eight published diploid potato genomes from samples of A6-26, E4-63, PG1008, PG1011, PG1013, PG3003, PG3005, and PG3022^32, 33^. All genome assemblies were downloaded from the database at http://solomics.agis.org.cn/potato/. The PacBio HiFi and Illumina sequencing reads of these were downloaded from the NCBI database, under accession number PRJNA641265 and PRJNA754534. The genome data from two autotetraploid cultivars, C88 and Otava, were downloaded from http://spuddb.uga.edu/. The Illumina and PacBio HiFi sequencing data for Otava were downloaded from the NCBI database under accession number PRJNA751899^20^. The Illumina and PacBio HiFi sequencing data for C88 were downloaded from the NGDC database under accession number CRA006012^21^.

#### Wheat

Genome assemblies of 10 cultivars were published in a previous study including ArinaLrFor, CDC_Landmark, CDC_Stanley, Jagger, Julius, LongReach_Lancer, Mace, Norin61, spelta_PI190962, SY_Mattis^34^. The corresponding Illumina sequencing reads were downloaded from the NCBI database under accession number PRJNA544491. The reference genome was IWGSC-v2.1^35^.

#### Human

Human population variant data were from the HPRC project^17^. Illumina sequencing reads of three samples (HG00438, HG01928, NA20129) were used for genotyping and downloaded from the NCBI database, under accession numbers PRJNA701308 and PRJNA731524.

### Simulated datasets

For *A. thaliana* and rice, we used simulated short reads for pangenome graph based variant genotyping. These reads were produced from genomes simulated based on random selection of real variants derived from assembly comparisons between the reference genome and several alternative genomes. We used minimap2 (version 2.24-r1122)^36^ to align each alternative genome to the reference genome with the parameters “-ax asm5 -eqx”, followed by calling different types of variations (SNPs, indels, and SVs) with the tool SyRI (version 1.6.3)^37^ under parameters of “-k -F s”. Finally, VarSim (version 0.8.6)^38^ was used to incorporate these variations into the reference genome to produce simulated genome with known variations.

Specifically, the parameters for simulating the *A. thaliana* genome were: -sv_num_ins 5000 --sv_num_del 5000 --sv_num_dup 2500 --sv_num_inv 2500 --vc_num_snp 500000 -- vc_num_ins 20000 --vc_num_del 20000 --vc_num_mnp 5000 --vc_num_complex 2500 -- vc_min_length_lim 0 --vc_max_length_lim 49 --sv_min_length_lim 50 --sv_max_length_lim 1000000 --disable_sim --vc_prop_het 0 --sv_prop_het 0.

The parameters for simulating the *Oryza sativa* genome were: --sv_num_ins 2500 -- sv_num_del 2500 --sv_num_dup 2000 --sv_num_inv 2000 --vc_num_snp 500000 -- vc_num_ins 10000 --vc_num_del 10000 --vc_num_mnp 2500 --vc_num_complex 2500 -- vc_min_length_lim 0 --vc_max_length_lim 49 --sv_min_length_lim 50 --sv_max_length_lim 1000000 --disable_sim --vc_prop_het 0.

In total, 65 *A. thaliana* alternative genomes and 251 *O. sativa* alternative genomes were generated. Next, for each species, variations from alternative genomes were merged for construction of a variant dataset and a pangenome graph. Besides, variants were then randomly selected from the dataset to simulate an additional genome to be genotyped.

Specifically, the parameters for the *A. thaliana* genome were: --sv_num_ins 5000 -- sv_num_del 5000 --sv_num_dup 2500 --sv_num_inv 2500 --vc_num_snp 500000 -- vc_num_ins 20000 --vc_num_del 20000 --vc_num_mnp 5000 --vc_num_complex 2500 -- vc_min_length_lim 0 --vc_max_length_lim 49 --sv_min_length_lim 50 --sv_max_length_lim 1000000 --disable_sim --vc_prop_het 0 --sv_prop_het 0. For the rice genome, the parameters were: --sv_num_ins 10000 --sv_num_del 10000 --sv_num_dup 2500 --sv_num_inv 2500 -- vc_num_snp 2000000 --vc_num_ins 40000 --vc_num_del 40000 --vc_num_mnp 5000 -- vc_num_complex 2500 --vc_min_length_lim 0 --vc_max_length_lim 49 --sv_min_length_lim 50 --sv_max_length_lim 1000000 --disable_sim --vc_prop_het 0 --sv_prop_het 0.

Based on this additional genome, second-generation sequencing reads were simulated using ART (version 2.18) with the parameters “-s 10 -p -l 150 -m 600”. These paired-end short reads were used for variant genotyping for each pangenome graph-based genotyper.

Besides, to evaluate the performance of genotypers on heterozygous genomes, we simulated 66 heterozygous diploid *A. thaliana* genomes using VarSim with parameters of “-- vc_prop_het 0.6” and “--sv_prop_het 0.6” based on the same variation dataset as above. For one genome, we also used ART to simulate short reads for variant genotyping. Additionally, we produced 33 autotetraploid *A. thaliana* genomes by random recombination of these 66 heterozygous genomes. We used BCFtools merge (version 1.9)^39^ to merge variant files (VCF format) from multiple samples to create a population variant set. Again, we also used ART (version 2.1.8)^40^ to simulate short reads for variant genotyping for one genome.

### Benchmarking variants from real datasets

To construct high-quality real variant datasets for benchmarking, we developed a pipeline by integrating evidence from comparisons of genome assemblies and alignments of sequencing reads from multiple platforms and employing different algorithms of variant calling. In principle, this pipeline leverage advantages of each data type, such as the high accuracy of second-generation sequencing data, the long-read length of third-generation sequencing, the high-quality of *de novo* genome assemblies, ensuring the accuracy of these variant datasets. This construction strategy has been successfully applied to build human benchmark variant set in GIAB and HGSV^41, 42^. Here, this pipeline was used to construct real variant datasets for maize, pummelo, rapeseed, and potato.

The construction pipeline comprises three parts: genome assembly-based, long reads-based, and short reads-based variant detection **(Supplementary Fig. 1)**.

#### Genome assembly-based variant detection

First, nucmer from the MUMmer4 package (version 4.0.0rc1)^43^ was used to align each alternative genome assembly to the reference genome with the parameters “--maxmatch -c 100 -b 500 -l 50”. Next, the alignments were filtered using delta-filter^43^, retaining only those with an alignment rate greater than 90%, alignment length greater than 10,000 bp, and the most optimal alignment, with parameters “-i 90 -l 10000 -1”. The filtered delta files were then converted to coordinate files using show-coords^43^, followed by identifying SNPs and indels using delta2vcf^43^. Additionally, SyRI was used to detect SNPs, indels, and SVs with default parameters, and Assemblytics (version 1.2.1)^44^ was used to identify SVs (retaining only variants between 50 bp and 1,000,000 bp in length).

#### Long read-based variant detection

First, minimap2 was used to align the long-read data to the reference genome with the parameter “-eqx”. SVs were identified using cuteSV (version 2.1.0, parameters: --max_cluster_bias_INS 1000 --diff_ratio_merging_INS 0.9 -- max_cluster_bias_DEL 1000 --diff_ratio_merging_DEL 0.5 --genotype), DeBreak (version 1.3, parameters: --rescue_large_ins --rescue_dup --poa), Sniffles (version 2.2, default parameters), and SVIM (version 2.0.0, default parameters)^45–48^. If the sequencing data were from PacBio HiFi, DeepVariant (version 1.6.0)^49^ was used to call SNPs and indels from the alignment, with the parameter “--model_type=PACBIO”. Only variants with a quality score (QUAL) ≥ 10 and depth (DP) ≥ 5 were retained to filter out low-quality sites.

#### Short read-based variant detection

BWA-MEM (version 0.7.17-r1198-dirty)^50^ was used to align short reads to the reference genome, followed by sorting and converting the alignments into BAM format using SAMtools (version 1.15-8)^39^. SNPs and indels were then identified using DeepVariant and GATK (version 4.5.0.0)^23^, respectively. Variants with a quality score (QUAL) ≥ 10 and depth (DP) ≥ 5 were retained to filter out low-quality sites.

The identified variants from different methods described as above were merged using jasmine (version 1.1.5)^51^, retaining only variants supported by two or more tools, with the parameters “max_dist=1000 min_support=2 --dup_to_ins --ignore_type --ignore_strand -- normalize_type”. Finally, a high-quality variant set was generated for each species, to construct pan-genome graphs.

#### Variant dataset for autotetraploid potato

Chromosome-level and haplotype-resolved assemblies from two autotetraploid potato cultivars, C88 and Otava, are available, enabling precise variant identification and genotyping^20, 21^. Apart from GATK, other sequencing read-based variant calling tools are only suitable for diploid genomes. However, the raw Illumina and PacBio HiFi data are from mixed haplotypes. To run these tools, we separated the sequencing reads into each haplotype before variant calling. Specifically, we first aligned the Illumina and PacBio HiFi reads to the genome assembly using BWA-MEM and minimap2, respectively. Reads were then assigned to the haplotype with the best alignment. If one read were aligned to multiple haplotypes with equal alignment scores, it was randomly assigned to one of the haplotypes. The effectiveness of haplotype phasing was evaluated by examining the *k*-mers depth distribution of the phasing data **(Supplementary Fig. 17)**. Then, the phased reads and each haplotype assembly were used to call variants for each haplotype based on the pipeline as described above. For each cultivar, variants from each haplotype were then merged to construct a variant dataset.

Finally, for each of these species, maize, pummelo, rapeseed, and potato, the variant from different samples were merged to generate a population-level variant dataset using BCFtools^39^. This dataset was used for pangenome graph construction and variant genotyping.

### The Varigraph algorithm

In brief, Varigraph performs variant genotyping without doing direct read mapping, but mainly based on the comparisons of *k*-mers at variant sites and *k*-mers of short reads from a given genome to be genotyped. It first constructs a variant graph based on the reference genome and variants from alternative genomes. Then, using reference genome and genome graph, Varigraph builds a Counting Bloom Filter and the *k*-mer index. Next, Varigraph scans the sequencing reads from an individual to be genotyped and counts the depth of *k*-mers in the genome graph index. Finally, Varigraph utilizes the *k*-mer depth information in the genome graph to calculate the posterior probabilities of all candidate genotypes at each variant site based on the Hidden Markov Model. Varigraph can output the genotypes, posterior probabilities, and genotype quality at all variant sites. The detailed algorithm is described in the following subsections.

#### Construction of pangenome graphs

Given a reference genome and population variant VCF files, Varigraph incorporates all variant into the coordinates of a reference genome and constructs a Directed Acyclic Graph (DAG) of variations. Each variant in the graph forms a “bubble”, containing multiple nodes that represent different haplotypes at that site. Nodes correspond to the sample haplotype across different bubbles and are connected by edges. For example, haplotype 1 (hap1) in bubble 1 will be connected to hap1 in bubble 2.

During the construction of the genome graph, Varigraph checks if the haplotype 0 sequence (reference genome, corresponding to the REF sequence in the VCF file) at each bubble matches the corresponding sequence in the reference genome. If there is a discrepancy, the reference genome sequence is used to replace the REF sequences in the VCF. For the ALT sequences in the VCF file, as Varigraph can directly verify their corresponding sequences in the alternative genome, the ALT sequences are inserted into the graph directly. This requires that ALT sequences in the given VCF file do not contain special characters.

#### Index of the genome graph

The Varigraph indexing consists of two main components: 1) a Counting Bloom Filter for the reference genome, and 2) a *k*-mer index for the genome graph.

##### Construction of a Counting Bloom Filter

Varigraph uses a Counting Bloom Filter instead of a hash table to store the frequency information of all *k*-mers in the reference genome. If the Bloom Filter uses *k* hash functions, the time complexity of query is *O(k)*. Although it is slightly slower than a hash table, the time complexity remains constant, making it an efficient data structure with memory usage approximately ten times the size of the genome. The false positive rate of the Bloom Filter can be calculated using Equation 1. By taking the derivative of its approximation and finding the minimum value, the optimal number of hash functions, *k*, can be determined, given that the Bloom Filter size is *m* and the expected number of elements is *n*, as shown in Equation 2.

Once the Bloom Filter is initialized, Varigraph splits the reference genome into *k*-mers and generates *k* hash values for each *k*-mer using the efficient non-cryptographic hash function MurmurHash3. The corresponding positions in the Bloom Filter are incremented by 1. The Bloom Filter created by Varigraph uses 8-bit numbers, so the maximum frequency for each *k*-mer is capped at 255.

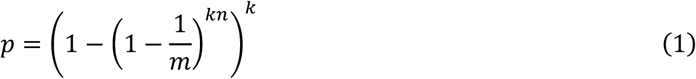

Where *p* represents the false positive rate of the Bloom Filter, *m* is the size of the Bloom Filter, *n* is the expected number of inserted elements, and *k* is the number of hash functions.

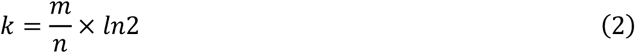

Where *k* is the number of hash functions, *m* is the size of the Bloom Filter, and *n* is the expected number of inserted elements.

##### Construction of the genome graph k-mer index

Based on the constructed variant graph, Varigraph splits the sequences of each node within every bubble into *k*-mers. Since some node sequences may be shorted than the *k*-mer length, varigraph extends the sequence of each node by *k-1* bp upstream and downstream before constructing the index. This ensures that even if a node represents a SNP, the extended sequence will still contain *k k*-mers. If the extension process encounters another bubble, the sequence from the corresponding haplotype within the bubble is used for filling.

K-mers that appear more than once in the genome can skew the depth distribution in sequencing data. Therefore, Varigraph only load k-mers with a frequency of less than 2 (node-specific *k*-mers) from the Counting Bloom Filter. However, when bubbles occur in repetitive sequence regions, or the variation itself is a repetitive sequence, a node may not contain any unique *k*-mers. In such cases, Varigraph selects the minimum frequency of all *k*-mers from all nodes in the bubble as the threshold to ensure that every bubble contains characteristic *k*-mers.

Assume that each *k*-mer corresponds to a structure *S*{*d*, *f*, *h*}, where *d* represents the depth of the *k*-mer in the sequencing reads, *f* represents the frequency of the k-mer in the genome graph, and *h* contains information on which haplotypes include that k-mer. Instead of using a list to store *h*, Varigraph adopts a bitmap approach. Specifically, multiple 8-bit numbers are used to represent the haplotype information. If the number of haplotypes in the VCF file is *N*, then ⌈*N/*8⌉ numbers are used to store *h*. The initial 8-bit number is set to 00000000. If haplotype 1 contains the *k*-mer, the binary representation is updated to 00000001. The number 3 (00000011) indicates that both haplotypes 1 and 2 contain the *k*-mer. Using this bitmap data structure, Varigraph reduces memory consumption by 16-fold. For example, if a k-mer’s structure is *S*{21, 2, 3}, it indicates that the *k*-mer’s depth in the sequencing reads is 21, it appears twice in the graph (with an actual depth of 21/2), and haplotype 1 and 2 both contain the *k*-mer.

#### Counting k-mer depth in sequencing data

After constructing the genome graph index, Varigraph processes the sequencing data by splitting the reads into k-mers. Assuming that the function used to traverse the sequencing reads is *read_count()*, the initial return value is the set of all *k*-mers from the given read. To reduce memory copying and movement, Varigraph only returns *k*-mers that appear in the genome graph index and increments the count *d* in the corresponding *k*-mer structure in the graph index by 1.

#### Varigraph’s genotyping model

Once the depth, frequency, and haplotype information of all *k*-mers in the genome graph have been obtained, Varigraph performs genotype inference using a Hidden Markov Model (HMM):

##### Hidden states

Assume the genome graph contains *N* haplotypes (*H_i_*, where *i* ∈ {1, 2, …, *N*}) and *M* bubbles (*B_j_*, where j ∈ {1, 2, …, *M*}). Each bubble corresponds to a list of *k*-mers (*K*), where *K* contains *L k*-mers, and *K*[*i*] (where *l* ∈ {1, 2, …, *L*}) represents the *l*-th *k*-mer in the list. This *k*-mer points to a structure *S*{*d*, *f*, *h*}. As described in previous section, *S.d* represents the depth of *k*-mer in the sequencing data, *S.f* represents its frequency in the genome graph, and *S.h* represents its haplotype information.

For any bubble *B_j_*, the hidden state *η_v_* is defined as the set of all possible haplotype combinations: {*H_v_*_, *i*, *j*, *k*, *l*_ | *i*, *j*, *k*, *l* ≤ *N*}. Each hidden state contains the cope number of every *k*-mer list *K* under the current haplotype combination. For tetraploid genomes, the cope number can range from 0 to 4, where 0 indicates that none of the haplotypes in the current combination contains the *k*-mer, and 1, 2, 3, and 4 represent the presence of 1, 2, 3, 4 copies of the *k*-mer in the haplotype combination, respectively. The specific formula is as follows:

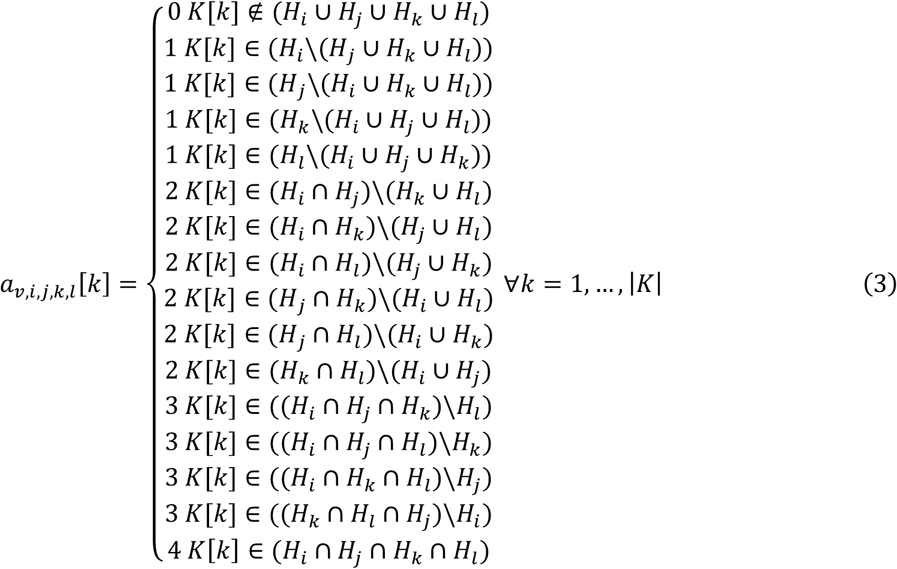

Thus, the length of the hidden state list (*η_v_*) is the number of all possible haplotype combinations, with each hidden state corresponding to a list of copy numbers for each *k*-mer (*a_v_*_,*i*,*j*,*k*,*l*_). Here, *a_v,i,j,k,l_*[*k*] represents the copy number of the *k*-th *k*-mer under the current haplotype combination.

##### Transition probability

Let *kmerD* denote the total depth of all *k*-mers for all haplotype in the genome graph, and *kmerD*[*i*] represent the total depth of haplotype *i*. Varigraph calculates the probability of each haplotype appearing based on a Sparse Dirichlet Distribution. The shape parameter of the Dirichlet distribution is set to *kmerD*[*i*] + 1.0, and the scale parameter is 1. The specific formula is as follows:

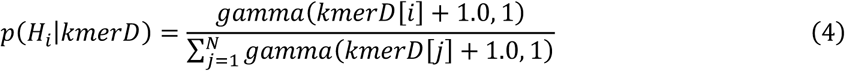

*p*(*H_i_*│*kmerD*) represents the probability of haplotype *i* appearing in the population given the current *kmerD* depth.

##### Observation matrix

For tetraploid genomes, at any given bubble, each hidden state in *η_v_* outputs the copy number of each *k*-mer under the current haplotype combination (0, 1, 2, 3, or 4). Let the haplotype *k*-mer depth in the sequencing reads be represented as *λ*. If the copy number of a *k*-mer is 4, a Poisson distribution with a mean of 4 × *λ* is used to calculate the probability of that *k*-mer when the actual depth is *S.d*. Similarly, if the copy number is 3, a Poisson distribution with a mean of 3 × *λ* is used, and the same applies for copy numbers of 2 and 1. When the copy number is 0, it is assumed that the *k*-mer is a sequencing error. The specific formula is as follows:

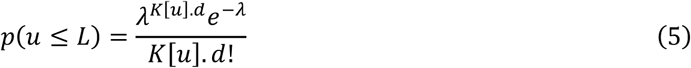

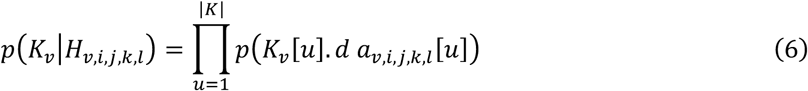

Equation 5 defines the Poisson distribution, where *u* ranges from 1 to *L*. *K*[*u*].*d* represents the depth of the *u*-th *k*-mer in the *k*-mers list, and *λ* is the expected depth of the k-mer.

In equation 6, *p*(*K_v_*│*H_v,i,j,k,l_*) denotes the probability that the *k*-mers at bubble *v* have depth *K_v_* given the haplotype combination *H_i,j,k,l_* at that bubble.

##### Variant genotyping

The genotyping algorithm is similar to PanGenie, utilizing the Forward-Backward Hidden Markov Model (HMM) to compute the probability of each haplotype combination.

##### Forward algorithm

For bubble 1 through *M*, the initial probability is set to 1 (Equation 7). The forward algorithm is computed as follows:

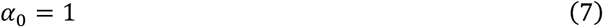

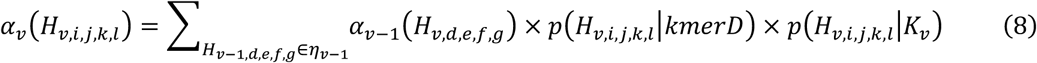

##### Backward algorithm

For bubble *M* to 1, the initial probability is set to 1 (Equation 9). The backward algorithm is computed as follows:

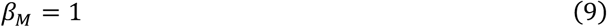

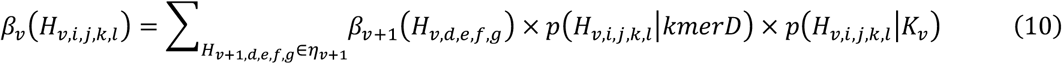

##### Posterior probability

Finally, based on the results of the forward and backward algorithms, the posterior probability for each haplotype combination at each bubble is computed using the following formula:

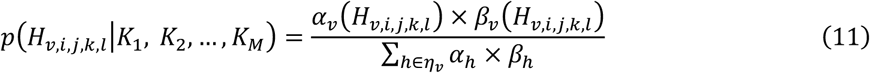

### Comparison with other genotyping software

The genotyping tool and their respective versions used in this study are as follows: varigraph (version 1.0.7), vg (version 1.50.1)^3, 5^, BayesTyper (version 1.5)^12^, GraphTyper2 (version 2.7.2)^7^, Paragraph (version 2.3)^2^, PanGenie (version 3.0.1)^8^, GATK (version 4.5.0.0)^23^, FreeBayes (version 1.3.6)^52^, and Octopus (version 0.7.4)^53^.

Due to the limitation that PanGenie supports a maximum of 254 haplotypes, it was tested using variant information from only 127 samples in the rice dataset, while other software utilized variant information from all samples.

The parameters used for each software are as follows:

- varigraph: --use-depth --granularity 100
- vg: default parameters
- GraphAligner: default parameters
- BayesTyper: -z -p 10 --noise-genotyping
- GraphTyper2: --avg_cov_by_readlen
- Paragraph: --scratch-dir
- PanGenie: -g
- GATK: --sample-ploidy 4
- FreeBayes: -p 4
- Octopus: -P 4

The server configuration used for testing was an Intel(R) Xeon(R) Platinum 8375C @ 2.90GHz CPU and NVIDIA GeForce RTX 4090 GPU. All software was tested using 10 threads.

## Data availability

All real genome assemblies, variants, and sequencing reads used for evaluation were accessed from public databases, except the data of pummelo genomes and sequencing reads. The accession IDs of these datasets are shown in Methods above. The genome assemblies, and PacBio HiFi long reads of pummelo have been deposited into National Genomics Data Center (BioProject ID PRJCA035310) and China National GeneBank DataBase (BioProject ID CNP0006728).

## Code availability

The Varigraph software is released under the MIT license and is freely available on GitHub: https://github.com/JiaoLab2021/varigraph. The command lines used in this study are provided in GitHub: https://github.com/JiaoLab2021/varigraph.

## Declarations

### Ethics approval and consent to participate

Not applicable.

### Consent for publication

Not applicable.

### Authors’ contributions

WBJ conceived and designed the project. ZZD and JBH collected the data. ZZD and JBH conducted the analyses. ZZD developed the Varigraph. ZZD and WBJ wrote the paper with contribution from PXX, JH, NY. All authors read and approved the final manuscript.

## Acknowledgements

We would like to thank Jeffrey Ross-Ibarra (University of California, Davis) for helpful discussions.

