## Supplementary Figures and Tables for "Varigraph: an accurate and widely applicable pangenome graph-based variant genotyper for diploid and polyploid genomes"

### Supplementary Information

### Supplementary Figures

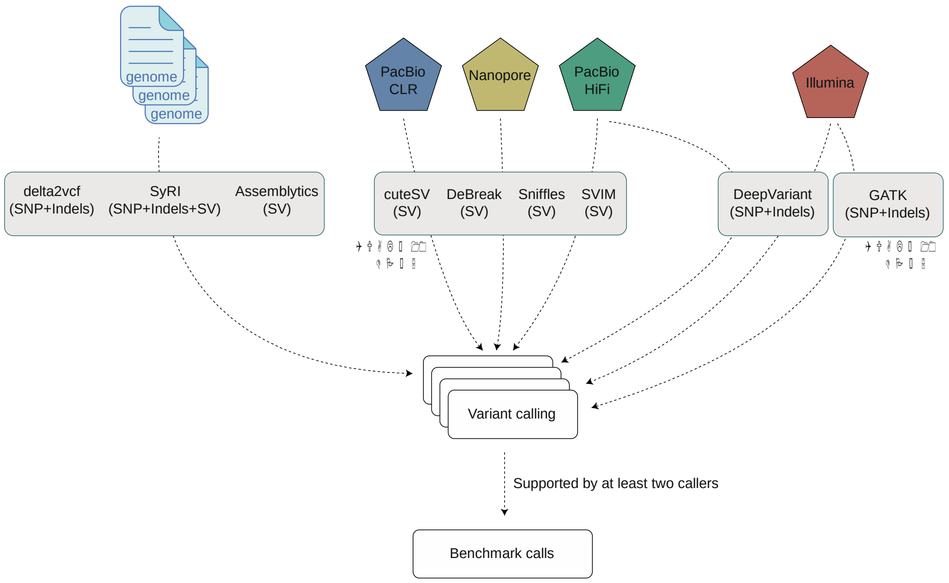


**Supplementary Fig. 1. The workflow of benchmark variant calling.** According to the type of datasets, the workflow includes three parts: **1)** identify all kinds of variants based on genome assemblies, **2)** detect variants using third-generation long-reads, **3)** identify SNPs and indels using second-generation short reads. For variants identified through second- and third-generation sequencing, only variants with quality score (QUAL) ≥10 and depth (DP) ≥5 were retained to filter out low-quality ones.

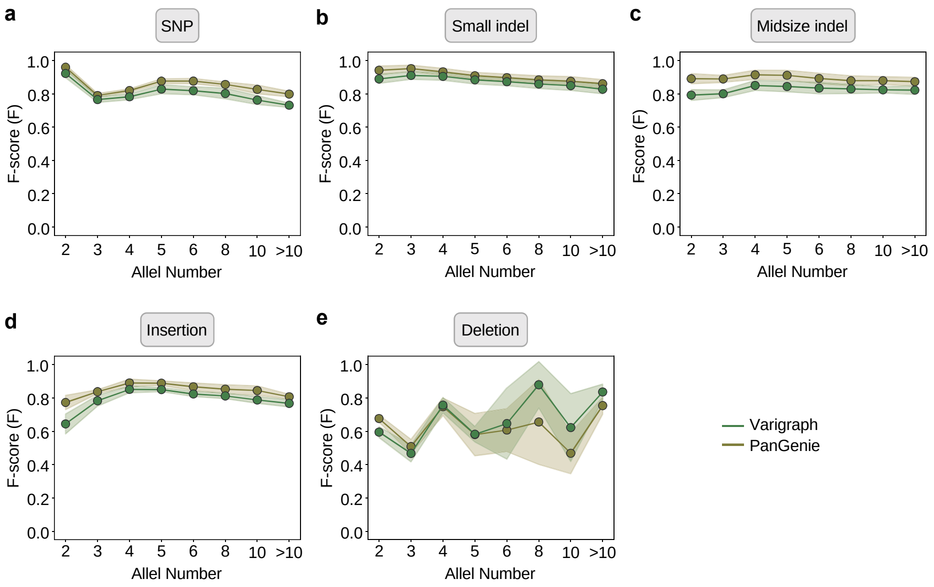


**Supplementary Fig. 2. Genotyping performance on bi-allelic and multi-allelic variants of the HPRC dataset of human genomes.** Small indel:1-19 bp, Midsize indel: 20-49 bp. Insertion and Deletion: >=50 bp.

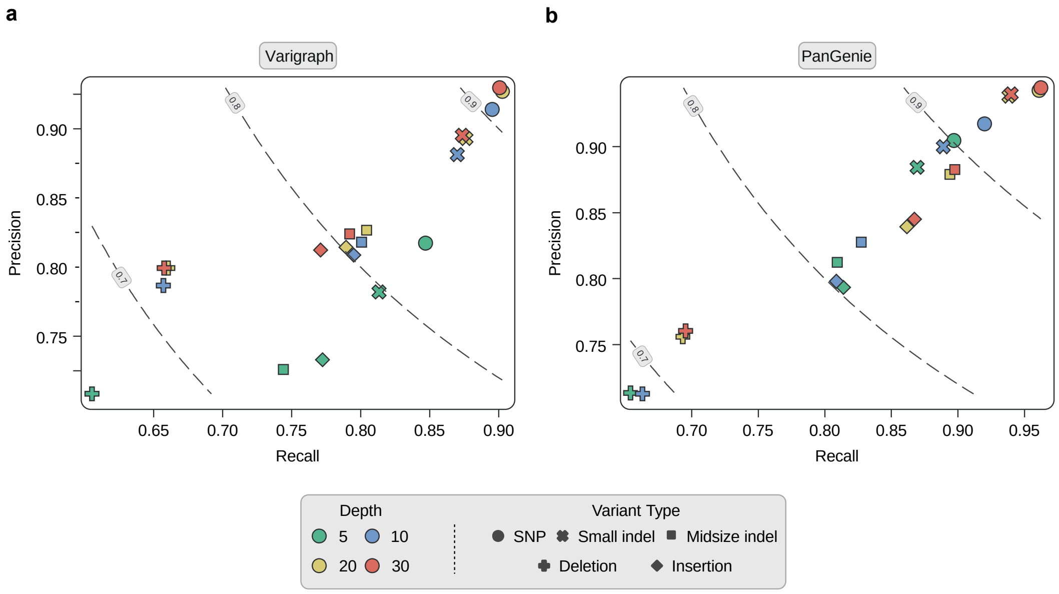


**Supplementary Fig. 3. Genotyping performance of varigraph and PanGenie under different sequencing depths based on the HPRC dataset of human genomes.** Sequencing depths in the figures include five gradients: 5×, 10×, 20×, and 30×.

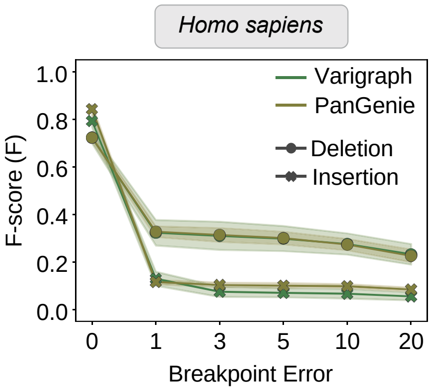


**Supplementary Fig. 4. Genotyping performance of varigraph and PanGenie under different breakpoint errors based on the HPRC dataset of human genomes.** Different size of breakpoint deviations to the real variants were introduced.

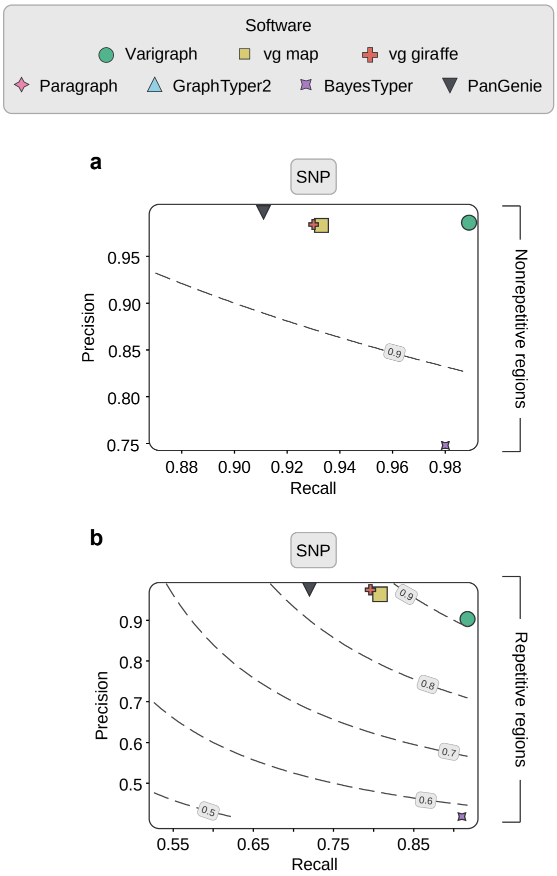


**Supplementary Fig. 5. Performance of different pangenome graph-based variant genotypers on based on simulated data of *A. thaliana* genomes.** The dashed lines correspond to the recall and precision when F-scores vary. Numbers on the dashed lines indicate F-scores. The genome graph was constructed based on one reference genome and 66 alternative genomes. Paired-end (2 × 150 bp) short reads with 30× depth were simulated for genotyping. Paragraph and GraphTyper2 were used only for SV genotyping due to the computational limitation of huge number of SNPs and indels.

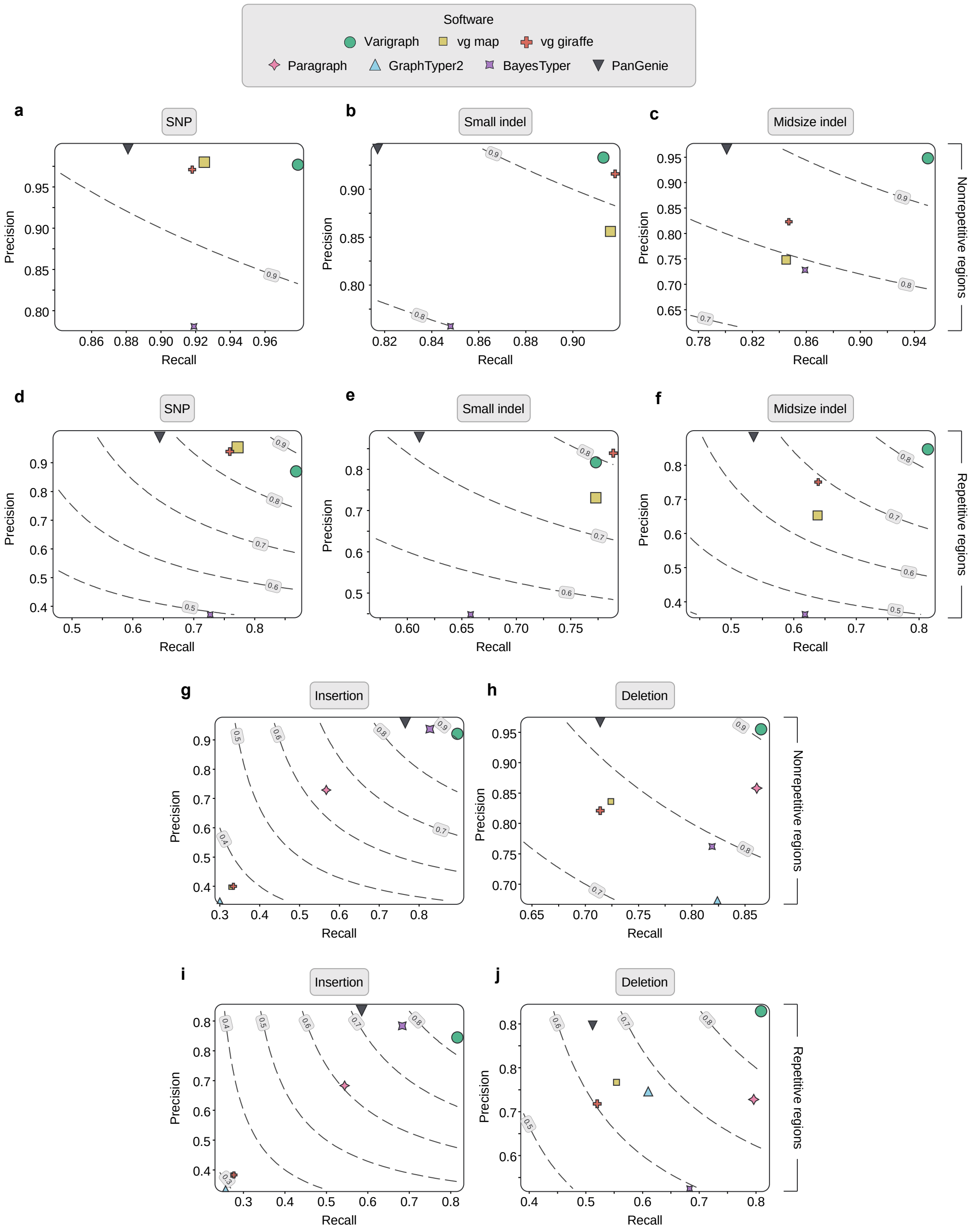


**Supplementary Fig. 6. Performance of different pangenome graph-based variant genotypers based on simulated data of synthetic heterozygous diploid *A. thaliana* genomes.** Genotyping performance was visualized for SNP, small indel (1-19 bp), midsize indel (20-49 bp), insertion (>=50 bp), and deletions (>=50 bp) in nonrepetitive **(a-c, g-h)** and repetitive regions **(d-f, i-j)** of *A. thaliana* genomes. The dashed lines correspond to recall and precision under different F-scores. Numbers on the dashed lines indicate F-scores. The genome graph was constructed based on one reference genome and 66 synthetic heterozygous genomes derived by introducing know variants into the reference genome. Paired-end (2 × 150 bp) short reads with 30× depth were simulated for genotyping. Paragraph and GraphTyper2 were used only for SV genotyping due to the computational limitation of huge number of SNPs and indels.

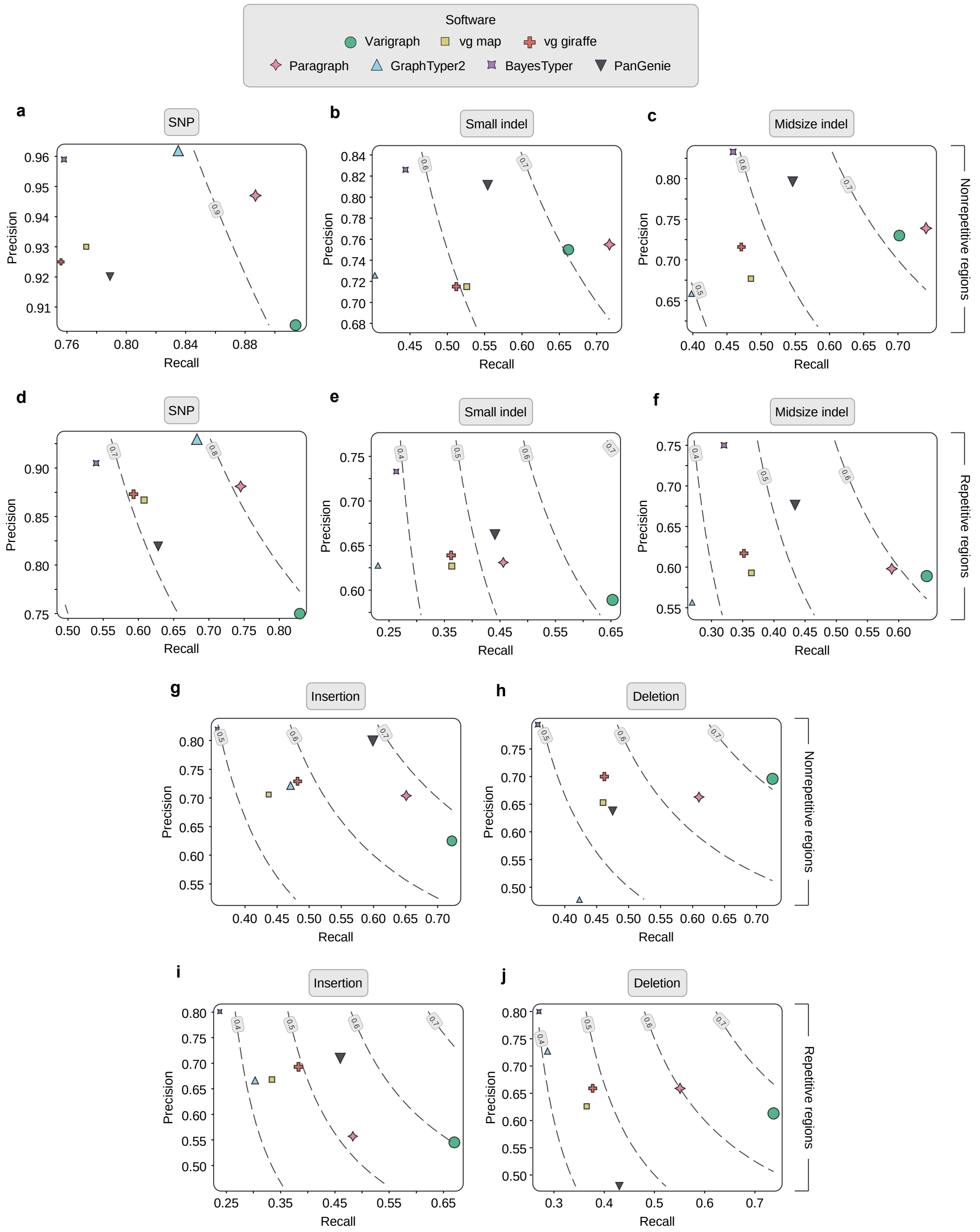


**Supplementary Fig. 7. Performance of different pangenome graph-based variant genotypers based on real data of heterozygous diploid *Citrus maxima* genomes.** Genotyping performance was visualized for small indel (1-19 bp), midsize indel (20-49 bp), insertion (>=50 bp), and deletions (>=50 bp) in nonrepetitive **(a-c, g-h)** and repetitive regions **(d-f, i-j)** of *C. maxima* genomes. The dashed lines correspond to recall and precision under different F-scores. Numbers on the dashed lines indicate F-scores. The genome graph was constructed based on one reference genome and 7 alternative genomes. Paired-end (2 × 123 bp) short reads from the accession GBY with 28.8× depth were used for genotyping. Notably, a haplotype-resolved assembly and original PacBio HiFi long reads from accession GBY were used for constructing the benchmarking dataset of sequence variants.

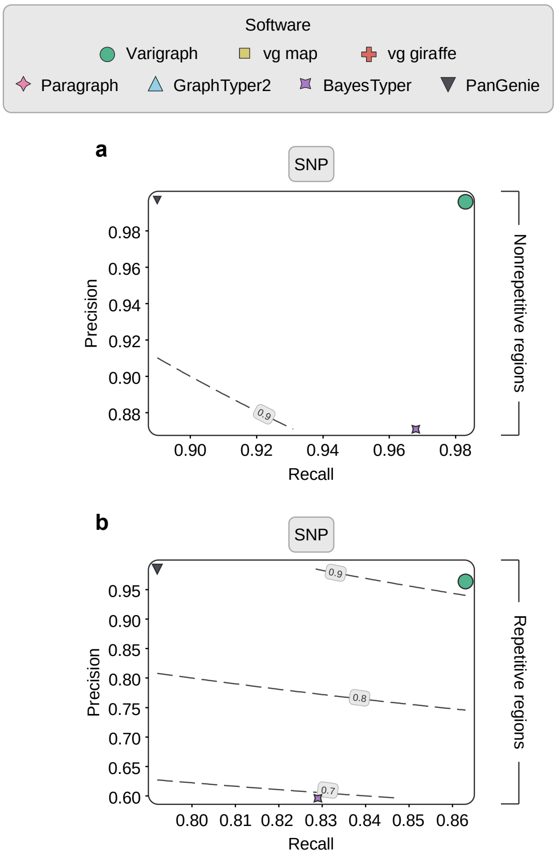


**Supplementary Fig. 8. Genotyping performance of SNPs between varigraph and other pangenome graph-based variant genotypers for a large pangenome graph with 252 rice genomes.** The dashed lines correspond to recall and precision under different F-scores. Numbers on the dashed lines indicate F-scores.

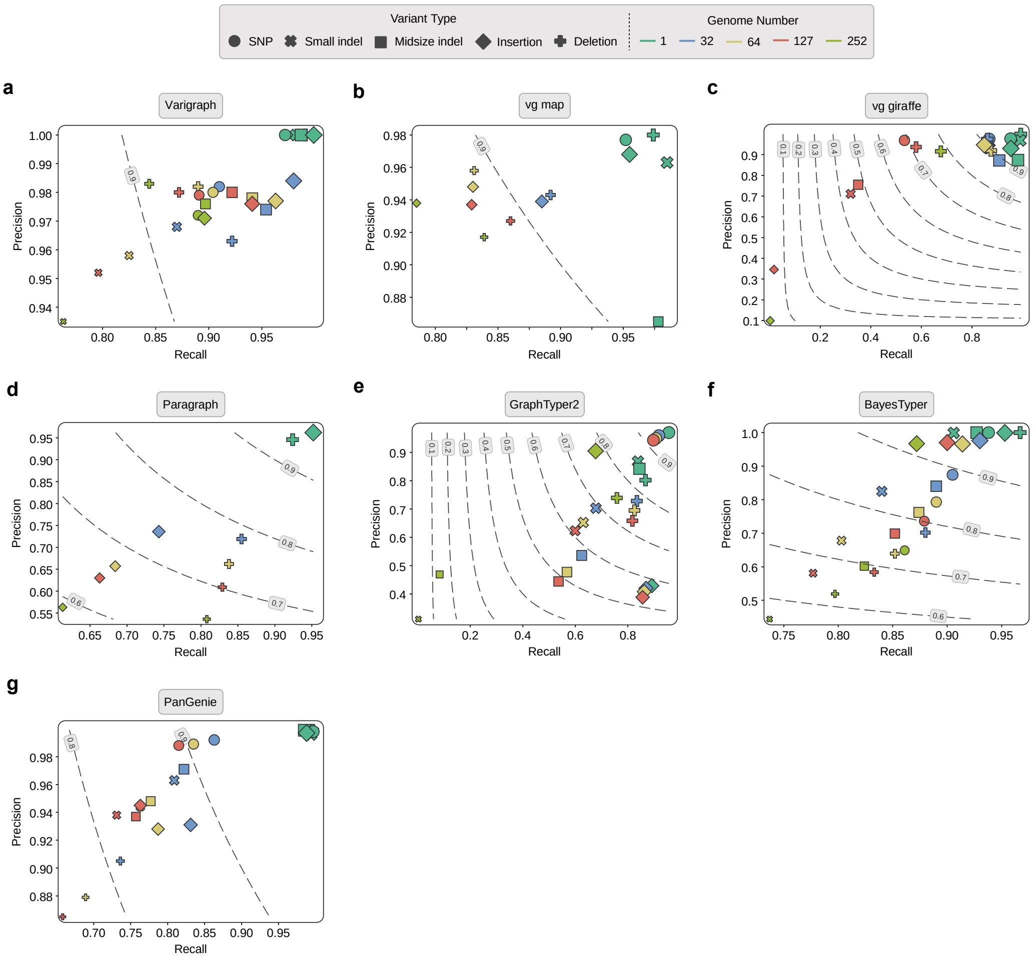


**Supplementary Fig. 9. Genotyping performance of pangenome graph based variant genotypers across graphs with different number of rice genomes.** Genotyping performance for SNP, small indel (1-19 bp), midsize indel (20-49 bp), insertion (>=50 bp), and deletions (>=50 bp). The dashed lines correspond to recall and precision under different F-scores. Numbers on the dashed lines indicate F-scores. The maximum genome number for testing PanGenie was 127 as it can only handle up to 127 diploid genomes.

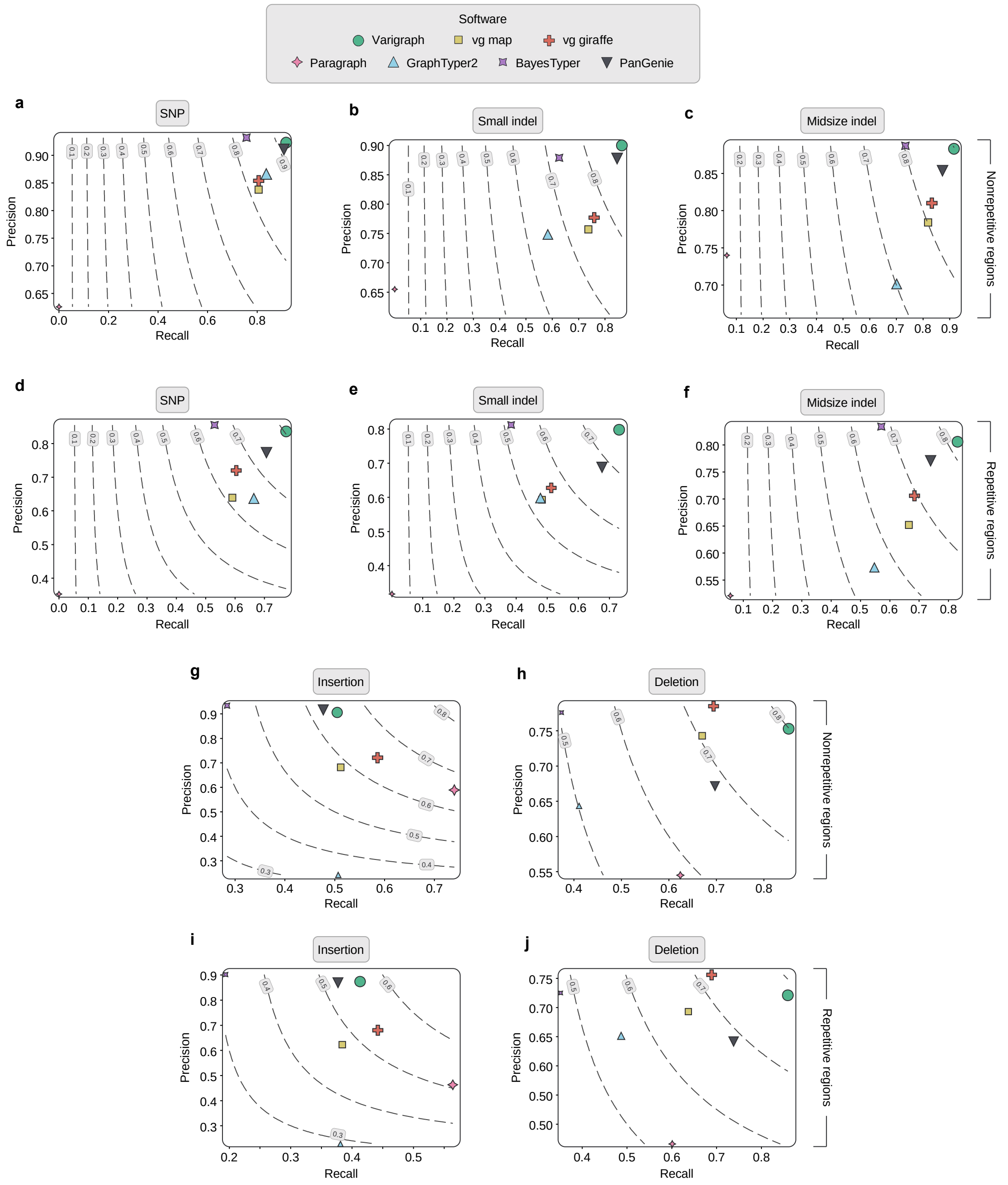


**Supplementary Fig. 10. Genotyping performance of graph based variant genotypers based on real dataset of *Brassica napus* genomes.** Genotyping performance for SNP, small indel (1-19 bp), midsize indel (20-49 bp), insertion (>=50 bp), and deletions (>=50 bp) in nonrepetitive **(a-c, g-h)** and repetitive regions **(d-f, i-j)**. The dashed lines correspond to recall and precision under different F-scores. Numbers on the dashed lines indicate F-scores. The genome graph was constructed based on one reference genome and seven homozygous alternative genomes. Paired-end (2 × 147 bp) short reads (Westar) with 116.3× depth are used for genotyping.

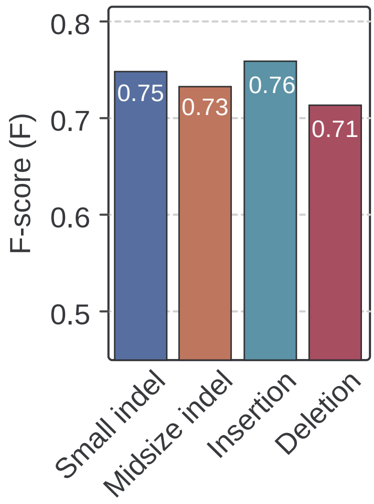


**Supplementary Fig. 11. Genotyping performance of varigraph based on real dataset of bread wheat (*Triticum aestivum*) genomes.** Small indel: 1-19 bp, midsize indel: 20-49 bp, insertion: >=50 bp, deletions: >=50 bp. The genome graph was constructed based on one reference genome and 10 homozygous alternative genomes. Paired-end (2 × 149 bp) short reads from the accession Mace with 44.2× depth were used for genotyping.

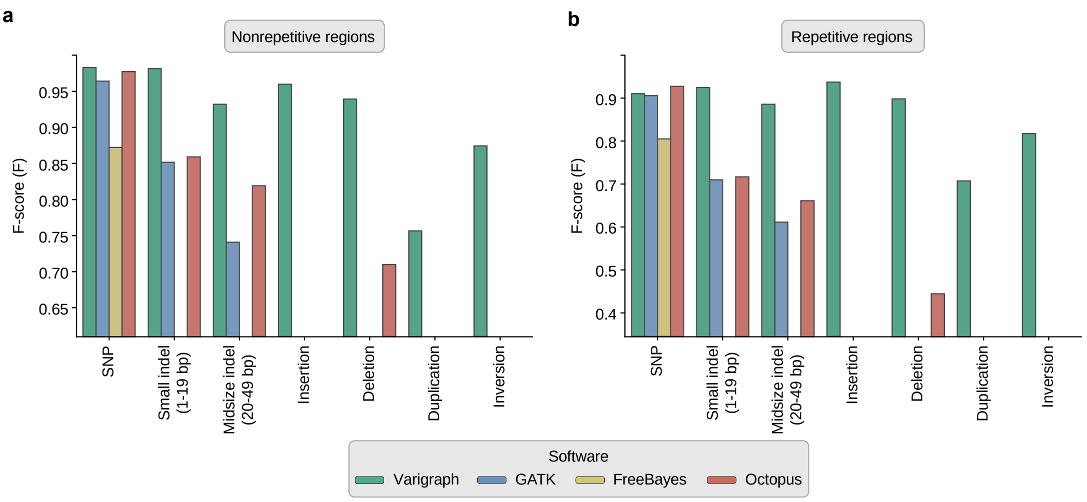


**Supplementary Fig. 12. Genotyping performance of Varigraph and GATK on simulated autotetraploid *A. thaliana* genomes.** The genome graph was constructed based on one reference genome and 33 homozygous tetraploid alternative genomes. Paired-end (2 × 150 bp) short reads (Westar) with 60× depth are simulated for genotyping.

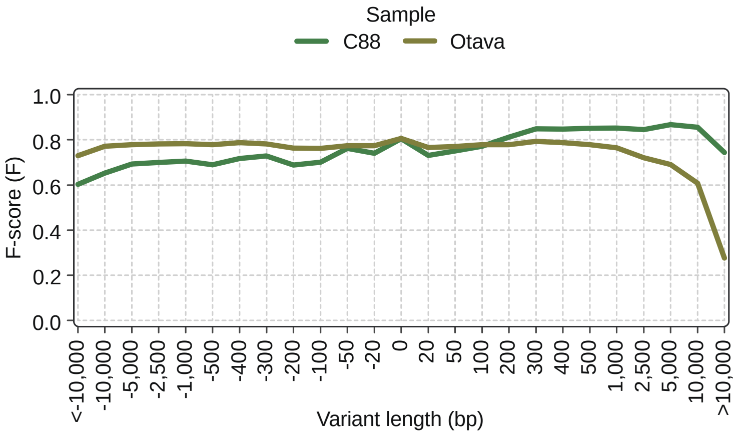


**Supplementary Fig. 13. Genotyping performance of insertions and deletions with vary length in autotetraploid potato genomes.**

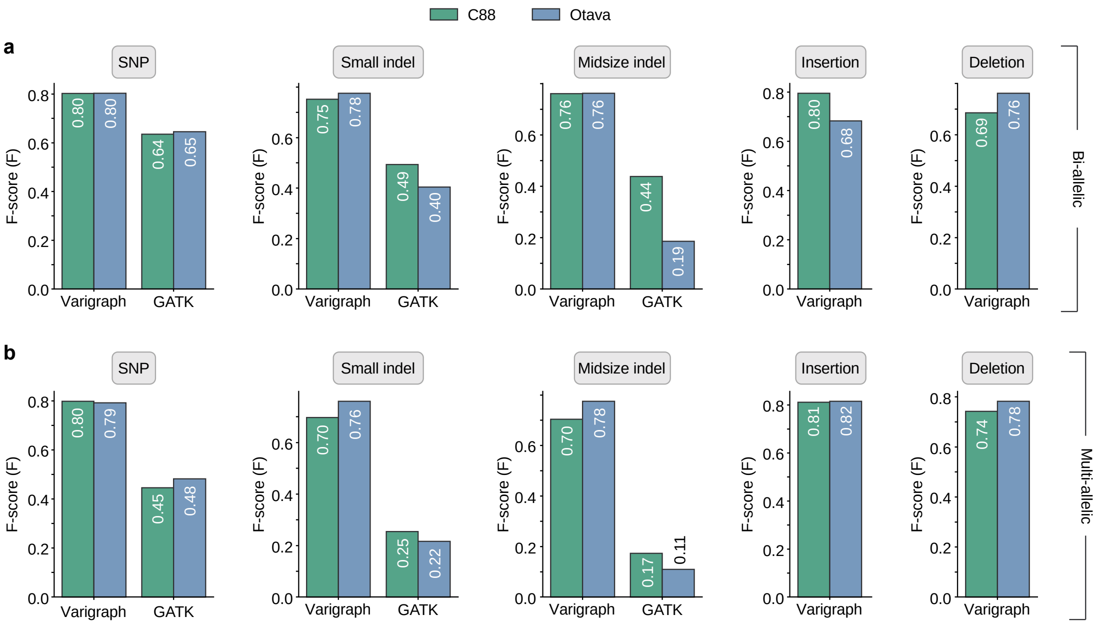


**Supplementary Fig. 14. Genotyping F-scores of Varigraph and GATK on bi-allelic and multi-allelic variants based on real data (*Solanum tuberosum*).** Small indel: 1-19 bp, midsize indel: 20-49 bp, insertion: >=50 bp, deletions: >=50 bp.

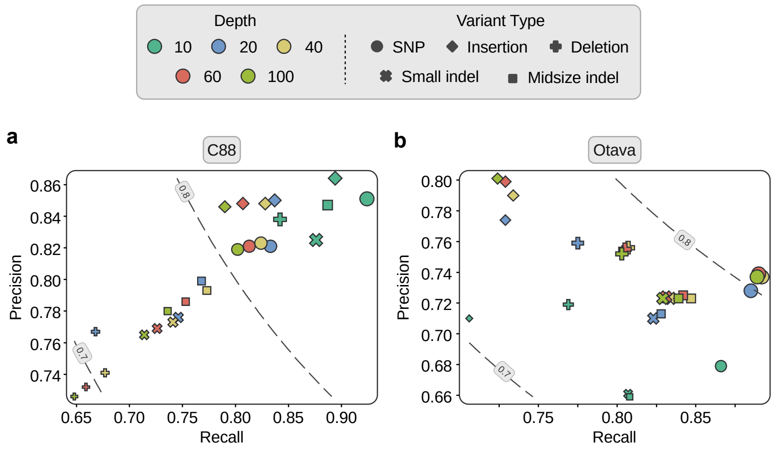


**Supplementary Fig. 15. Genotyping performance of Varigraph under different sequencing depths based on real data (*Solanum tuberosum*).** The dashed lines correspond to recall and precision under different F-scores. Numbers on the dashed lines indicate F-scores. The genome graph was constructed using one reference genome and 10 tetraploid alternative genomes. Sequencing depths in the figures include five gradients: 10×, 20×, 40×, 60×, and 100×.

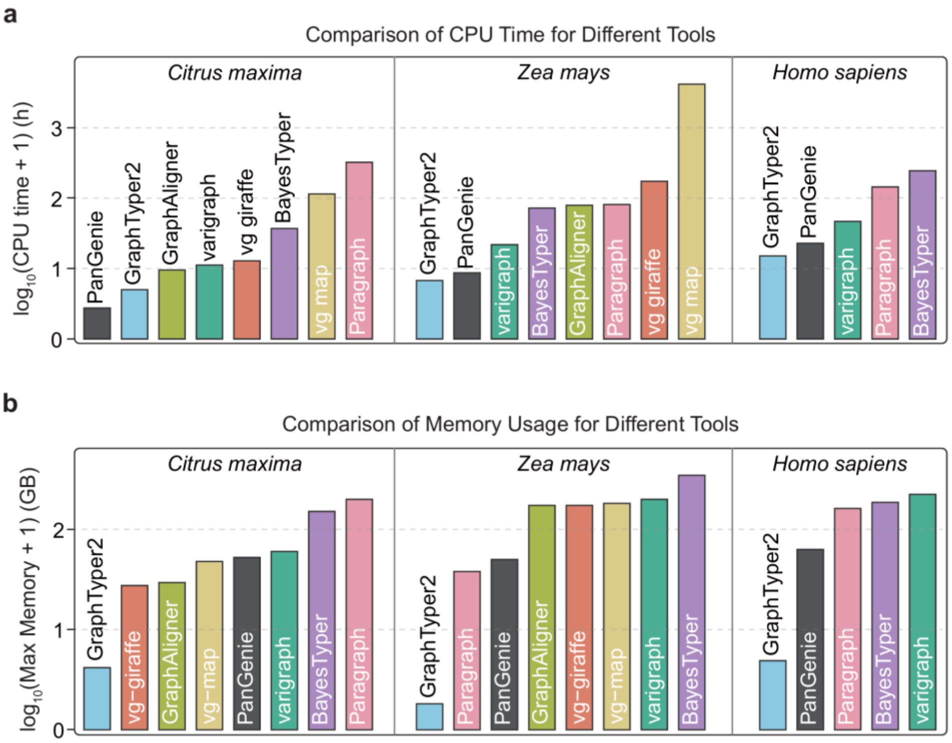


**Supplementary Fig. 16. Runtime ana memory usage for genotyper in different plant genomes.** The server's CPU is Intel(R) Xeon(R) Platinum 8375C @ 2.90GHz. All software tests were conducted using 10 threads.

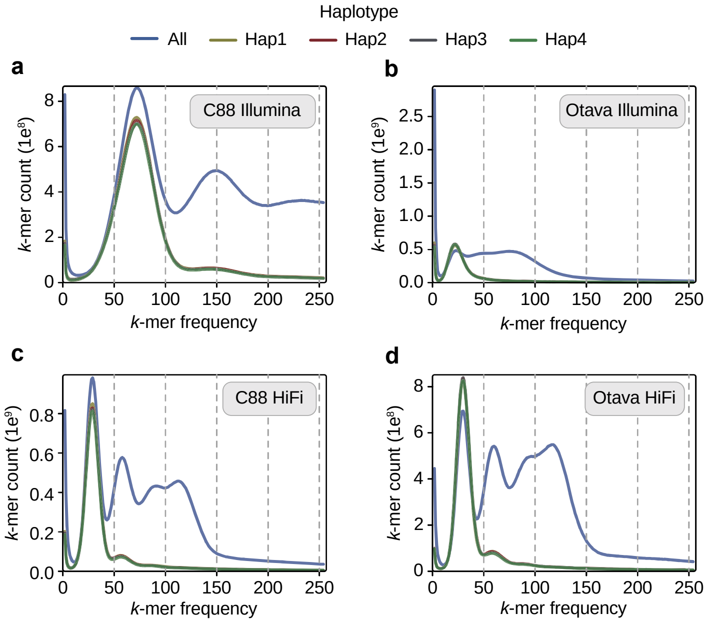


**Supplementary Fig. 17. *k*-mer frequency distribution of split tetraploid potato data. (a)** and **(b)** display the Illumina data split results for C88 and Otava, respectively. **(c)** and **(d)** show the PacBio HiFi data split results for C88 and Otava, respectively.

#### Supplementary Tables

**Supplementary Table 1. Comparisons of Varigraph running time on CPU and GPU.**

| Genome | Counting Bloom Filter | | Read (k-mer) | |
|---|---|---|---|---|
| | CPU (s) | GPU (s) | CPU (s) | GPU (s) |
| *Arabidopsis thaliana* | 47 | 1 | 414 | 351 |
| *Oryza sativa* | 442 | 46 | 1389 | 500 |
| *Citrus maxima* | 417 | 42 | 1134 | 430 |
| *Solanum tuberosum* | 294 | 87 | 10271 | 4350 |
| *Brassica napus* | 569 | 111 | 12153 | 3795 |
| *Zea mays* | 1271 | 285 | 1490 | 2389 |

Note: The CPU used was an Intel(R) Xeon(R) Platinum 8375C @ 2.90GHz and the GPU was an NVIDIA GeForce RTX 4090.

**Supplementary Table 2. Summary of reference genomes used in this study.**

| Organism | Karyotype | Accession | Genome size (Mb) | Repeat Content (%) |
|---|---|---|---:|---:|
| *Arabidopsis thaliana* | 2n = 2x = 10 | Col-0 | 119 | 21.4 |
| *Oryza sativa* | 2n = 2x = 24 | Nipponbare | 373 | 58 |
| *Citrus maxima* | 2n = 2x = 18 | CMJ | 345 | 57.36 |
| *Brassica napus* | 2n = 4x = 38 | ZS11 | 961 | 65.6 |
| *Zea mays* | 2n = 2x = 20 | B73 | 2,132 | 88.9 |
| *Solanum tuberosum* | 2n = 2x = 24 | DMv8 | 738 | 71.34 |
| *Triticum aestivum* | 2n = 6x = 42 | IWGSC | 14,066 | 85 |
| *Homo sapiens* | 2n = 2x = 46 | GRCh38 | 3,100 | 51.1 |

**Supplementary Table 3. Distribution of variants across size ranges in pangenome graphs with varying numbers of genomes**

See the file Supplementary.Table.3.xlsx

**Supplementary Table 4. Cumulative size distribution of variants across size ranges in pangenome graphs with varying numbers of genomes.**

See the file Supplementary.Table.4.xlsx

**Supplementary Table 5. Summary of simulated short reads for variant genotyping.**

| Organism | Homozygous/Heterozygous | Genome ploidy | Read length (bp) | Depth (×) | Read number (bp) | Read base (bp) |
|---|---|---|---|---|---|---|
| *A. thaliana* | Homozygous | 2 | 150 | 30 | 24,100,656 | 3,615,098,400 |
| *A. thaliana* | Homozygous | 2 | 150 | 5 | 3,993,344 | 599,001,600 |
| *A. thaliana* | Homozygous | 2 | 150 | 10 | 7,986,612 | 1,197,991,800 |
| *A. thaliana* | Homozygous | 2 | 150 | 20 | 15,973,174 | 2,395,976,100 |
| *A. thaliana* | Homozygous | 2 | 150 | 30 | 23,959,912 | 3,593,986,800 |
| *A. thaliana* | Homozygous | 2 | 150 | 50 | 39,933,160 | 5,989,974,000 |
| *A. thaliana* | Heterozygous | 4 | 150 | 60 | 47,770,760 | 7,165,614,000 |
| *O. sativa* | Homozygous | 2 | 150 | 30 | 74,864,726 | 11,229,708,900 |

**Supplementary Table 6. Summary of real genome assemblies, long-read, and short-read datasets used in this study.**

See the Supplementary.Table.6.xlsx

**Supplementary Table 7. The corresponding recall and precision of variant genotyping in a human pangenome graph for Fig. 2.**

See the file Supplementary.Table.7.xlsx.

**Supplementary Table 8. The corresponding recall and precision of variant genotyping in a A. thaliana pangenome graph for Fig. 3a-h.**

| Regions | Variation type | Metrics | Varigraph | vg map | vg giraffe | Paragraph | BayesTyper | GraphTyper2 | PanGenie |
|---|---|---|---|---|---|---|---|---|---|
| Nonrepetitive | SNPs | Recall | 0.99 | 0.93 | 0.93 | - | 0.98 | - | 0.91 |
| | | Precision | 0.99 | 0.98 | 0.98 | - | 0.75 | - | 1.00 |
| | | F-score | 0.99 | 0.96 | 0.96 | - | 0.85 | - | 0.95 |
| | Small indel | Recall | 0.94 | 0.94 | 0.95 | - | 0.91 | - | 0.86 |
| | | Precision | 0.95 | 0.88 | 0.94 | - | 0.73 | - | 0.95 |
| | | F-score | 0.94 | 0.91 | 0.94 | - | 0.81 | - | 0.90 |
| | Midsize indel | Recall | 0.98 | 0.92 | 0.92 | - | 0.91 | - | 0.84 |
| | | Precision | 0.96 | 0.81 | 0.90 | - | 0.67 | - | 0.95 |
| | | F-score | 0.97 | 0.86 | 0.91 | - | 0.77 | - | 0.89 |
| | Deletion | Recall | 0.92 | 0.89 | 0.89 | 0.88 | 0.90 | 0.89 | 0.77 |
| | | Precision | 0.98 | 0.98 | 0.99 | 0.86 | 0.74 | 0.73 | 0.95 |
| | | F-score | 0.95 | 0.93 | 0.94 | 0.87 | 0.81 | 0.80 | 0.85 |
| | Insertion | Recall | 0.96 | 0.84 | 0.84 | 0.58 | 0.94 | 0.78 | 0.81 |
| | | Precision | 1.00 | 0.96 | 0.96 | 0.58 | 0.99 | 0.99 | 0.98 |
| | | F-score | 0.98 | 0.89 | 0.90 | 0.58 | 0.96 | 0.87 | 0.89 |
| Repetitive | SNPs | Recall | 0.92 | 0.81 | 0.80 | - | 0.91 | - | 0.72 |
| | | Precision | 0.90 | 0.96 | 0.98 | - | 0.42 | - | 0.99 |
| | | F-score | 0.91 | 0.88 | 0.88 | - | 0.57 | - | 0.83 |
| | Small indel | Recall | 0.82 | 0.82 | 0.83 | - | 0.78 | - | 0.65 |
| | | Precision | 0.84 | 0.76 | 0.87 | - | 0.46 | - | 0.88 |
| | | F-score | 0.83 | 0.79 | 0.85 | - | 0.58 | - | 0.75 |
| | Midsize indel | Recall | 0.84 | 0.76 | 0.77 | - | 0.82 | - | 0.62 |
| | | Precision | 0.84 | 0.69 | 0.83 | - | 0.35 | - | 0.90 |
| | | F-score | 0.84 | 0.72 | 0.80 | - | 0.49 | - | 0.74 |
| | Deletion | Recall | 0.86 | 0.77 | 0.77 | 0.81 | 0.83 | 0.70 | 0.62 |
| | | Precision | 0.95 | 0.96 | 0.98 | 0.70 | 0.53 | 0.82 | 0.88 |
| | | F-score | 0.91 | 0.85 | 0.86 | 0.75 | 0.65 | 0.76 | 0.73 |
| | Insertion | Recall | 0.92 | 0.70 | 0.71 | 0.55 | 0.87 | 0.69 | 0.65 |
| | | Precision | 0.98 | 0.92 | 0.94 | 0.55 | 0.97 | 0.97 | 0.95 |
| | | F-score | 0.95 | 0.80 | 0.81 | 0.55 | 0.92 | 0.80 | 0.77 |

**Supplementary Table 9. Genotyping recall, precision, and F-measure of graph-based genotypers for variants in synthetic heterozygous *A. thaliana* genomes**

| Regions | Variation type | Metrics | Varigraph | vg map | vg giraffe | Paragraph | BayesTyper | GraphTyper2 | PanGenie |
|---|---|---|---|---|---|---|---|---|---|
| Nonrepetitive | SNPs | Recall | 0.98 | 0.93 | 0.92 | - | 0.92 | - | 0.88 |
| | | Precision | 0.98 | 0.98 | 0.97 | - | 0.78 | - | 1.00 |
| | | F-score | 0.98 | 0.95 | 0.94 | - | 0.84 | - | 0.93 |
| | Small indel | Recall | 0.91 | 0.92 | 0.92 | - | 0.85 | - | 0.82 |
| | | Precision | 0.93 | 0.86 | 0.92 | - | 0.76 | - | 0.94 |
| | | F-score | 0.92 | 0.88 | 0.92 | - | 0.80 | - | 0.88 |
| | Midsize indel | Recall | 0.95 | 0.85 | 0.85 | - | 0.86 | - | 0.80 |
| | | Precision | 0.95 | 0.75 | 0.82 | - | 0.73 | - | 0.97 |
| | | F-score | 0.95 | 0.79 | 0.83 | - | 0.79 | - | 0.88 |
| | Deletion | Recall | 0.87 | 0.72 | 0.71 | 0.86 | 0.82 | 0.82 | 0.71 |
| | | Precision | 0.96 | 0.84 | 0.82 | 0.86 | 0.76 | 0.67 | 0.97 |
| | | F-score | 0.91 | 0.78 | 0.76 | 0.86 | 0.79 | 0.74 | 0.82 |
| | Insertion | Recall | 0.90 | 0.33 | 0.33 | 0.57 | 0.83 | 0.30 | 0.77 |
| | | Precision | 0.92 | 0.40 | 0.40 | 0.73 | 0.94 | 0.35 | 0.96 |
| | | F-score | 0.91 | 0.36 | 0.36 | 0.64 | 0.88 | 0.32 | 0.85 |
| Repetitive | SNPs | Recall | 0.87 | 0.77 | 0.76 | - | 0.73 | - | 0.64 |
| | | Precision | 0.87 | 0.95 | 0.94 | - | 0.37 | - | 0.99 |
| | | F-score | 0.87 | 0.85 | 0.84 | - | 0.49 | - | 0.78 |
| | Small indel | Recall | 0.77 | 0.77 | 0.79 | - | 0.66 | - | 0.61 |
| | | Precision | 0.82 | 0.73 | 0.84 | - | 0.45 | - | 0.88 |
| | | F-score | 0.79 | 0.75 | 0.81 | - | 0.53 | - | 0.72 |
| | Midsize indel | Recall | 0.81 | 0.64 | 0.64 | - | 0.62 | - | 0.54 |
| | | Precision | 0.85 | 0.65 | 0.75 | - | 0.36 | - | 0.88 |
| | | F-score | 0.83 | 0.65 | 0.69 | - | 0.46 | - | 0.67 |
| | Deletion | Recall | 0.81 | 0.55 | 0.52 | 0.80 | 0.68 | 0.61 | 0.51 |
| | | Precision | 0.93 | 0.77 | 0.72 | 0.73 | 0.53 | 0.75 | 0.90 |
| | | F-score | 0.86 | 0.64 | 0.60 | 0.76 | 0.59 | 0.67 | 0.65 |
| | Insertion | Recall | 0.82 | 0.28 | 0.28 | 0.54 | 0.68 | 0.26 | 0.59 |
| | | Precision | 0.85 | 0.38 | 0.38 | 0.68 | 0.88 | 0.34 | 0.93 |
| | | F-score | 0.83 | 0.32 | 0.32 | 0.61 | 0.77 | 0.29 | 0.72 |

**Supplementary Table 10. Genotyping recall, precision, and F-measure of graph-based genotypers for variants in pummelo genomes.**

| Regions | Variation type | Metrics | Varigraph | vg map | vg giraffe | Paragraph | BayesTyper | GraphTyper2 | PanGenie |
|---|---|---|---|---|---|---|---|---|---|
| Nonrepetitive | SNPs | Recall | 0.91 | 0.77 | 0.76 | 0.89 | 0.76 | 0.84 | 0.79 |
| | | Precision | 0.90 | 0.93 | 0.93 | 0.95 | 0.96 | 0.96 | 0.92 |
| | | F-score | 0.91 | 0.84 | 0.83 | 0.92 | 0.85 | 0.89 | 0.85 |
| | Small indel | Recall | 0.66 | 0.53 | 0.51 | 0.72 | 0.44 | 0.40 | 0.55 |
| | | Precision | 0.75 | 0.72 | 0.72 | 0.76 | 0.83 | 0.73 | 0.81 |
| | | F-score | 0.70 | 0.61 | 0.60 | 0.74 | 0.58 | 0.52 | 0.66 |
| | Midsize indel | Recall | 0.70 | 0.49 | 0.47 | 0.74 | 0.46 | 0.40 | 0.55 |
| | | Precision | 0.73 | 0.68 | 0.72 | 0.74 | 0.83 | 0.66 | 0.80 |
| | | F-score | 0.72 | 0.57 | 0.57 | 0.74 | 0.59 | 0.50 | 0.65 |
| | Deletion | Recall | 0.73 | 0.46 | 0.46 | 0.61 | 0.36 | 0.42 | 0.48 |
| | | Precision | 0.70 | 0.65 | 0.70 | 0.66 | 0.79 | 0.48 | 0.64 |
| | | F-score | 0.71 | 0.54 | 0.56 | 0.64 | 0.49 | 0.45 | 0.54 |
| | Insertion | Recall | 0.72 | 0.44 | 0.48 | 0.65 | 0.36 | 0.47 | 0.60 |
| | | Precision | 0.63 | 0.71 | 0.73 | 0.70 | 0.82 | 0.72 | 0.80 |
| | | F-score | 0.67 | 0.54 | 0.58 | 0.68 | 0.50 | 0.57 | 0.68 |
| Repetitive | SNPs | Recall | 0.83 | 0.61 | 0.59 | 0.75 | 0.54 | 0.68 | 0.63 |
| | | Precision | 0.75 | 0.87 | 0.87 | 0.88 | 0.91 | 0.93 | 0.82 |
| | | F-score | 0.79 | 0.71 | 0.71 | 0.81 | 0.68 | 0.79 | 0.71 |
| | Small indel | Recall | 0.65 | 0.36 | 0.36 | 0.46 | 0.26 | 0.23 | 0.44 |
| | | Precision | 0.59 | 0.63 | 0.64 | 0.63 | 0.73 | 0.63 | 0.66 |
| | | F-score | 0.62 | 0.46 | 0.46 | 0.53 | 0.39 | 0.34 | 0.53 |
| | Midsize indel | Recall | 0.65 | 0.36 | 0.35 | 0.59 | 0.32 | 0.27 | 0.43 |
| | | Precision | 0.59 | 0.59 | 0.62 | 0.60 | 0.75 | 0.56 | 0.68 |
| | | F-score | 0.62 | 0.45 | 0.45 | 0.59 | 0.45 | 0.36 | 0.53 |
| | Deletion | Recall | 0.74 | 0.37 | 0.38 | 0.55 | 0.27 | 0.29 | 0.43 |
| | | Precision | 0.61 | 0.63 | 0.66 | 0.66 | 0.80 | 0.73 | 0.48 |
| | | F-score | 0.67 | 0.46 | 0.48 | 0.60 | 0.40 | 0.41 | 0.45 |
| | Insertion | Recall | 0.67 | 0.33 | 0.38 | 0.48 | 0.24 | 0.30 | 0.46 |
| | | Precision | 0.55 | 0.67 | 0.69 | 0.56 | 0.80 | 0.67 | 0.71 |
| | | F-score | 0.60 | 0.45 | 0.49 | 0.52 | 0.37 | 0.42 | 0.56 |

**Supplementary Table 11. Genotyping recall, precision, and F-measure of graph-based genotypers for variants in maize genomes**

| Regions | Variation type | Metrics | Varigraph | vg map | vg giraffe | Paragraph | BayesTyper | GraphTyper2 | PanGenie |
|---|---|---|---|---|---|---|---|---|---|
| Nonrepetitive | Deletion | Recall | 0.78 | 0.68 | 0.67 | 0.60 | 0.11 | 0.17 | 0.71 |
| | | Precision | 0.69 | 0.68 | 0.69 | 0.27 | 0.61 | 0.21 | 0.29 |
| | | F-score | 0.73 | 0.68 | 0.68 | 0.37 | 0.18 | 0.19 | 0.41 |
| | Insertion | Recall | 0.81 | 0.46 | 0.29 | 0.97 | 0.32 | 0.31 | 0.55 |
| | | Precision | 0.95 | 0.17 | 0.11 | 0.10 | 0.86 | 0.11 | 0.75 |
| | | F-score | 0.87 | 0.25 | 0.15 | 0.17 | 0.46 | 0.16 | 0.63 |
| Repetitive | Deletion | Recall | 0.63 | 0.52 | 0.57 | 0.39 | 0.09 | 0.24 | 0.64 |
| | | Precision | 0.68 | 0.47 | 0.53 | 0.14 | 0.49 | 0.37 | 0.32 |
| | | F-score | 0.66 | 0.50 | 0.55 | 0.21 | 0.16 | 0.30 | 0.43 |
| | Insertion | Recall | 0.76 | 0.38 | 0.31 | 0.78 | 0.30 | 0.22 | 0.47 |
| | | Precision | 0.92 | 0.17 | 0.14 | 0.07 | 0.84 | 0.09 | 0.70 |
| | | F-score | 0.83 | 0.23 | 0.19 | 0.13 | 0.44 | 0.13 | 0.56 |

**Supplementary Table 12. Genotyping recall, precision, and F-measure of graph-based genotypers for variants in rice genomes**

| Regions | Variation type | Metrics | Varigraph | vg map | vg giraffe | Paragraph | BayesTyper | GraphTyper2 | PanGenie |
|---|---|---|---|---|---|---|---|---|---|
| Nonrepetitive | SNPs | Recall | 0.98 | - | - | - | 0.97 | - | 0.89 |
| | | Precision | 1.00 | - | - | - | 0.87 | - | 1.00 |
| | | F-score | 0.99 | - | - | - | 0.92 | - | 0.94 |
| | Small indel | Recall | 0.94 | - | - | - | 0.90 | - | 0.86 |
| | | Precision | 0.97 | - | - | - | 0.76 | - | 0.97 |
| | | F-score | 0.95 | - | - | - | 0.82 | - | 0.91 |
| | Midsize indel | Recall | 0.98 | - | - | - | 0.94 | - | 0.87 |
| | | Precision | 1.00 | - | - | - | 0.86 | - | 0.95 |
| | | F-score | 0.99 | - | - | - | 0.90 | - | 0.91 |
| | Deletion | Recall | 0.91 | 0.88 | 0.75 | 0.88 | 0.89 | 0.88 | 0.75 |
| | | Precision | 0.98 | 0.92 | 0.93 | 0.73 | 0.71 | 0.54 | 0.91 |
| | | F-score | 0.94 | 0.90 | 0.83 | 0.80 | 0.79 | 0.67 | 0.82 |
| | Insertion | Recall | 0.96 | 0.90 | 0.00 | 0.64 | 0.96 | 0.80 | 0.86 |
| | | Precision | 1.00 | 0.96 | 0.13 | 0.63 | 0.99 | 0.95 | 0.98 |
| | | F-score | 0.98 | 0.93 | 0.01 | 0.63 | 0.97 | 0.87 | 0.92 |
| Repetitive | SNPs | Recall | 0.86 | - | - | - | 0.83 | - | 0.79 |
| | | Precision | 0.96 | - | - | - | 0.60 | - | 0.99 |
| | | F-score | 0.91 | - | - | - | 0.69 | - | 0.88 |
| | Small indel | Recall | 0.71 | - | - | - | 0.68 | - | 0.69 |
| | | Precision | 0.92 | - | - | - | 0.38 | - | 0.93 |
| | | F-score | 0.80 | - | - | - | 0.49 | - | 0.79 |
| | Midsize indel | Recall | 0.88 | - | - | - | 0.79 | - | 0.73 |
| | | Precision | 0.97 | - | - | - | 0.55 | - | 0.93 |
| | | F-score | 0.92 | - | - | - | 0.65 | - | 0.82 |
| | Deletion | Recall | 0.84 | 0.83 | 0.67 | 0.80 | 0.79 | 0.71 | 0.65 |
| | | Precision | 0.98 | 0.92 | 0.91 | 0.52 | 0.50 | 0.74 | 0.86 |
| | | F-score | 0.90 | 0.87 | 0.77 | 0.63 | 0.61 | 0.72 | 0.74 |
| | Insertion | Recall | 0.87 | 0.75 | 0.00 | 0.61 | 0.84 | 0.64 | 0.73 |
| | | Precision | 0.96 | 0.93 | 0.09 | 0.54 | 0.96 | 0.88 | 0.93 |
| | | F-score | 0.92 | 0.83 | 0.00 | 0.57 | 0.90 | 0.74 | 0.82 |

**Supplementary Table 13. Genotyping precision, recall and F-scores of different graph-based genotypers across graphs with different number of alternative rice genomes (corresponding to Supplementary Fig. 10)**

| Number | Variation type | Metrics | Varigraph | vg map | vg giraffe | Paragraph | BayesTyper | GraphTyper2 | PanGenie |
|---|---|---|---|---|---|---|---|---|---|
| 1 | SNPs | Recall | 0.97 | 0.95 | 0.95 | - | 0.94 | 0.96 | 1.00 |
| | | Precision | 1.00 | 0.98 | 0.98 | - | 1.00 | 0.97 | 1.00 |
| | | F-score | 0.99 | 0.96 | 0.96 | - | 0.97 | 0.96 | 1.00 |
| | Small indel | Recall | 0.98 | 0.99 | 0.99 | - | 0.91 | 0.84 | 0.99 |
| | | Precision | 1.00 | 0.96 | 0.97 | - | 1.00 | 0.87 | 1.00 |
| | | F-score | 0.99 | 0.97 | 0.98 | - | 0.95 | 0.85 | 0.99 |
| | Midsize indel | Recall | 0.99 | 0.98 | 0.98 | - | 0.93 | 0.84 | 0.99 |
| | | Precision | 1.00 | 0.87 | 0.87 | - | 1.00 | 0.84 | 1.00 |
| | | F-score | 0.99 | 0.92 | 0.92 | - | 0.96 | 0.84 | 0.99 |
| | Deletion | Recall | 0.99 | 0.97 | 0.99 | 0.92 | 0.97 | 0.87 | 1.00 |
| | | Precision | 1.00 | 0.98 | 1.00 | 0.95 | 1.00 | 0.80 | 1.00 |
| | | F-score | 1.00 | 0.98 | 1.00 | 0.93 | 0.98 | 0.83 | 1.00 |
| | Insertion | Recall | 1.00 | 0.96 | 0.95 | 0.95 | 0.95 | 0.89 | 0.99 |
| | | Precision | 1.00 | 0.97 | 0.93 | 0.96 | 1.00 | 0.43 | 1.00 |
| | | F-score | 1.00 | 0.96 | 0.94 | 0.96 | 0.98 | 0.58 | 0.99 |
| 32 | SNPs | Recall | 0.91 | - | 0.87 | - | 0.91 | 0.92 | 0.86 |
| | | Precision | 0.98 | - | 0.98 | - | 0.87 | 0.96 | 0.99 |
| | | F-score | 0.94 | - | 0.92 | - | 0.89 | 0.94 | 0.92 |
| | Small indel | Recall | 0.87 | - | 0.87 | - | 0.84 | 0.68 | 0.81 |
| | | Precision | 0.97 | - | 0.93 | - | 0.83 | 0.70 | 0.96 |
| | | F-score | 0.92 | - | 0.90 | - | 0.83 | 0.69 | 0.88 |
| | Midsize indel | Recall | 0.95 | - | 0.91 | - | 0.89 | 0.62 | 0.82 |
| | | Precision | 0.97 | - | 0.87 | - | 0.84 | 0.54 | 0.97 |
| | | F-score | 0.96 | - | 0.89 | - | 0.86 | 0.58 | 0.89 |
| | Deletion | Recall | 0.92 | 0.89 | 0.86 | 0.86 | 0.88 | 0.83 | 0.74 |
| | | Precision | 0.96 | 0.94 | 0.98 | 0.72 | 0.70 | 0.73 | 0.91 |
| | | F-score | 0.94 | 0.92 | 0.91 | 0.78 | 0.78 | 0.78 | 0.81 |
| | Insertion | Recall | 0.98 | 0.89 | 0.86 | 0.74 | 0.93 | 0.87 | 0.83 |
| | | Precision | 0.98 | 0.94 | 0.95 | 0.74 | 0.98 | 0.42 | 0.93 |
| | | F-score | 0.98 | 0.91 | 0.90 | 0.74 | 0.95 | 0.57 | 0.88 |
| 64 | SNPs | Recall | 0.90 | - | - | - | 0.89 | 0.91 | 0.84 |
| | | Precision | 0.98 | - | - | - | 0.79 | 0.95 | 0.99 |
| | | F-score | 0.94 | - | - | - | 0.84 | 0.93 | 0.91 |
| | Small indel | Recall | 0.83 | - | - | - | 0.80 | 0.63 | 0.76 |
| | | Precision | 0.96 | - | - | - | 0.68 | 0.65 | 0.94 |

| | | | | | | | | | |
|---|---|---|---|---|---|---|---|---|---|
| | | F-score | 0.89 | - | - | - | 0.74 | 0.64 | 0.84 |
| | Midsize indel | Recall | 0.94 | - | - | - | 0.87 | 0.57 | 0.78 |
| | | Precision | 0.98 | - | - | - | 0.76 | 0.48 | 0.95 |
| | | F-score | 0.96 | - | - | - | 0.81 | 0.52 | 0.85 |
| | Deletion | Recall | 0.89 | 0.83 | 0.88 | 0.84 | 0.85 | 0.83 | 0.69 |
| | | Precision | 0.98 | 0.96 | 0.92 | 0.66 | 0.64 | 0.70 | 0.88 |
| | | F-score | 0.93 | 0.89 | 0.90 | 0.74 | 0.73 | 0.75 | 0.77 |
| | Insertion | Recall | 0.96 | 0.83 | 0.85 | 0.68 | 0.91 | 0.86 | 0.79 |
| | | Precision | 0.98 | 0.95 | 0.95 | 0.66 | 0.97 | 0.41 | 0.93 |
| | | F-score | 0.97 | 0.89 | 0.90 | 0.67 | 0.94 | 0.55 | 0.85 |
| 127 | SNPs | Recall | 0.89 | - | - | - | 0.88 | 0.90 | 0.82 |
| | | Precision | 0.98 | - | - | - | 0.74 | 0.94 | 0.99 |
| | | F-score | 0.93 | - | - | - | 0.80 | 0.92 | 0.89 |
| | Small indel | Recall | 0.80 | - | - | - | 0.78 | 0.60 | 0.73 |
| | | Precision | 0.95 | - | - | - | 0.58 | 0.62 | 0.94 |
| | | F-score | 0.87 | - | - | - | 0.66 | 0.61 | 0.82 |
| | Midsize indel | Recall | 0.92 | - | - | - | 0.85 | 0.54 | 0.76 |
| | | Precision | 0.98 | - | - | - | 0.70 | 0.44 | 0.94 |
| | | F-score | 0.95 | - | - | - | 0.77 | 0.49 | 0.84 |
| | Deletion | Recall | 0.87 | 0.86 | 0.58 | 0.83 | 0.83 | 0.82 | 0.66 |
| | | Precision | 0.98 | 0.93 | 0.94 | 0.61 | 0.58 | 0.66 | 0.87 |
| | | F-score | 0.92 | 0.89 | 0.72 | 0.70 | 0.69 | 0.73 | 0.75 |
| | Insertion | Recall | 0.94 | 0.83 | 0.02 | 0.66 | 0.90 | 0.86 | 0.76 |
| | | Precision | 0.98 | 0.94 | 0.35 | 0.63 | 0.97 | 0.39 | 0.95 |
| | | F-score | 0.96 | 0.88 | 0.03 | 0.65 | 0.93 | 0.53 | 0.84 |
| 252 | SNPs | Recall | 0.89 | - | - | - | 0.86 | - | - |
| | | Precision | 0.97 | - | - | - | 0.65 | - | - |
| | | F-score | 0.93 | - | - | - | 0.74 | - | - |
| | Small indel | Recall | 0.76 | - | - | - | 0.74 | - | - |
| | | Precision | 0.94 | - | - | - | 0.44 | - | - |
| | | F-score | 0.84 | - | - | - | 0.55 | - | - |
| | Midsize indel | Recall | 0.90 | - | - | - | 0.82 | - | - |
| | | Precision | 0.98 | - | - | - | 0.60 | - | - |
| | | F-score | 0.93 | - | - | - | 0.70 | - | - |
| | Deletion | Recall | 0.84 | 0.84 | 0.68 | 0.81 | 0.80 | 0.76 | - |
| | | Precision | 0.98 | 0.92 | 0.92 | 0.54 | 0.52 | 0.74 | - |
| | | F-score | 0.91 | 0.88 | 0.78 | 0.64 | 0.63 | 0.75 | - |
| | Insertion | Recall | 0.90 | 0.79 | 0.00 | 0.61 | 0.87 | 0.68 | - |
| | | Precision | 0.97 | 0.94 | 0.10 | 0.56 | 0.97 | 0.91 | - |
| | | F-score | 0.93 | 0.85 | 0.00 | 0.59 | 0.92 | 0.77 | - |

**Supplementary Table 14. Genotyping precision, recall and F-scores of graph based variant genotypers based on real dataset of *Brassica napus* genomes.**

| Regions | Variation type | Metrics | Varigraph | vg map | vg giraffe | Paragraph | BayesTyper | GraphTyper2 | PanGenie |
|---|---|---|---|---|---|---|---|---|---|
| Nonrepetitive | SNPs | Recall | 0.92 | 0.81 | 0.81 | - | 0.76 | 0.84 | 0.91 |
| | | Precision | 0.92 | 0.84 | 0.85 | - | 0.93 | 0.87 | 0.91 |
| | | F-score | 0.92 | 0.82 | 0.83 | - | 0.84 | 0.85 | 0.91 |
| | Small indel | Recall | 0.86 | 0.74 | 0.76 | - | 0.63 | 0.58 | 0.85 |
| | | Precision | 0.90 | 0.76 | 0.78 | - | 0.88 | 0.75 | 0.88 |
| | | F-score | 0.88 | 0.75 | 0.77 | - | 0.73 | 0.66 | 0.86 |
| | Midsize indel | Recall | 0.92 | 0.82 | 0.83 | - | 0.73 | 0.70 | 0.87 |
| | | Precision | 0.88 | 0.78 | 0.81 | - | 0.89 | 0.70 | 0.85 |
| | | F-score | 0.90 | 0.80 | 0.82 | - | 0.80 | 0.70 | 0.86 |
| | Deletion | Recall | 0.85 | 0.67 | 0.69 | 0.62 | 0.37 | 0.41 | 0.70 |
| | | Precision | 0.75 | 0.74 | 0.79 | 0.55 | 0.78 | 0.64 | 0.67 |
| | | F-score | 0.80 | 0.70 | 0.74 | 0.58 | 0.50 | 0.50 | 0.68 |
| | Insertion | Recall | 0.51 | 0.51 | 0.59 | 0.74 | 0.28 | 0.51 | 0.48 |
| | | Precision | 0.91 | 0.68 | 0.72 | 0.59 | 0.93 | 0.24 | 0.92 |
| | | F-score | 0.65 | 0.58 | 0.65 | 0.66 | 0.44 | 0.33 | 0.63 |
| Repetitive | SNPs | Recall | 0.77 | 0.59 | 0.60 | - | 0.53 | 0.66 | 0.71 |
| | | Precision | 0.84 | 0.64 | 0.72 | - | 0.86 | 0.64 | 0.77 |
| | | F-score | 0.80 | 0.61 | 0.66 | - | 0.65 | 0.65 | 0.74 |
| | Small indel | Recall | 0.73 | 0.48 | 0.51 | - | 0.38 | 0.48 | 0.68 |
| | | Precision | 0.80 | 0.59 | 0.63 | - | 0.81 | 0.60 | 0.69 |
| | | F-score | 0.76 | 0.53 | 0.56 | - | 0.52 | 0.53 | 0.68 |
| | Midsize indel | Recall | 0.83 | 0.67 | 0.68 | - | 0.57 | 0.55 | 0.74 |
| | | Precision | 0.81 | 0.65 | 0.71 | - | 0.83 | 0.57 | 0.77 |
| | | F-score | 0.82 | 0.66 | 0.69 | - | 0.68 | 0.56 | 0.75 |
| | Deletion | Recall | 0.86 | 0.64 | 0.69 | 0.60 | 0.35 | 0.49 | 0.74 |
| | | Precision | 0.72 | 0.69 | 0.76 | 0.47 | 0.73 | 0.65 | 0.64 |
| | | F-score | 0.78 | 0.66 | 0.72 | 0.52 | 0.47 | 0.56 | 0.69 |
| | Insertion | Recall | 0.41 | 0.38 | 0.44 | 0.56 | 0.19 | 0.38 | 0.38 |
| | | Precision | 0.87 | 0.62 | 0.68 | 0.46 | 0.90 | 0.23 | 0.87 |
| | | F-score | 0.56 | 0.48 | 0.54 | 0.51 | 0.32 | 0.29 | 0.53 |

**Supplementary Table 15. Genotyping precision, recall and F-scores of varigraph and GATK on simulated autotetraploid A. thaliana genomes.**

| Regions | Variation type | Metrics | Varigraph | GATK | FreeBayes | Octopus |
|---|---|---|---|---|---|---|
| Nonrepetitive | SNPs | Recall | 0.99 | 0.97 | 0.85 | 0.98 |
| | | Precision | 0.98 | 0.96 | 0.90 | 0.98 |
| | | F-score | 0.98 | 0.96 | 0.87 | 0.98 |
| | Small indel | Recall | 0.97 | 0.89 | - | 0.90 |
| | | Precision | 0.99 | 0.82 | - | 0.82 |
| | | F-score | 0.98 | 0.85 | - | 0.86 |
| | Midsize indel | Recall | 0.97 | 0.81 | - | 0.85 |
| | | Precision | 0.90 | 0.69 | - | 0.79 |
| | | F-score | 0.93 | 0.74 | - | 0.82 |
| | Deletion | Recall | 0.90 | - | - | 0.65 |
| | | Precision | 0.98 | - | - | 0.78 |
| | | F-score | 0.94 | - | - | 0.71 |
| | Insertion | Recall | 0.97 | - | - | - |
| | | Precision | 0.95 | - | - | - |
| | | F-score | 0.96 | - | - | - |
| Repetitive | SNPs | Recall | 0.89 | 0.91 | 0.80 | 0.92 |
| | | Precision | 0.94 | 0.90 | 0.81 | 0.93 |
| | | F-score | 0.91 | 0.91 | 0.80 | 0.93 |
| | Small indel | Recall | 0.90 | 0.77 | - | 0.78 |
| | | Precision | 0.96 | 0.66 | - | 0.67 |
| | | F-score | 0.92 | 0.71 | - | 0.72 |
| | Midsize indel | Recall | 0.88 | 0.69 | - | 0.68 |
| | | Precision | 0.90 | 0.55 | - | 0.64 |
| | | F-score | 0.89 | 0.61 | - | 0.66 |
| | Deletion | Recall | 0.84 | - | - | 0.32 |
| | | Precision | 0.97 | - | - | 0.72 |
| | | F-score | 0.90 | - | - | 0.44 |
| | Insertion | Recall | 0.94 | - | - | - |
| | | Precision | 0.94 | - | - | - |
| | | F-score | 0.94 | - | - | - |

**Supplementary Table 16. Genotyping precision, recall and F-scores of Varigraph and GATK on autotetraploid potato genomes. Corresponding to Figure 5.**

| Tools | Sample | Type | SNPs | | | Small indel | | | Midsize indel | | | Deletion | | | Insertion | | |
|---|---|---|---|---|---|---|---|---|---|---|---|---|---|---|---|---|---|
| | | | Recall | Precision | F-score | Recall | Precision | F-score | Recall | Precision | F-score | Recall | Precision | F-score | Recall | Precision | F-score |
| Varigraph | C88 | Nonrepetitive | 0.85 | 0.85 | 0.85 | 0.74 | 0.79 | 0.76 | 0.78 | 0.81 | 0.79 | 0.69 | 0.77 | 0.73 | 0.75 | 0.82 | 0.78 |
| | | Repetitive | 0.80 | 0.79 | 0.79 | 0.72 | 0.73 | 0.72 | 0.74 | 0.74 | 0.74 | 0.68 | 0.71 | 0.69 | 0.80 | 0.82 | 0.81 |
| | Otava | Nonrepetitive | 0.91 | 0.79 | 0.84 | 0.83 | 0.75 | 0.79 | 0.85 | 0.74 | 0.79 | 0.79 | 0.76 | 0.77 | 0.83 | 0.86 | 0.84 |
| | | Repetitive | 0.89 | 0.72 | 0.80 | 0.83 | 0.71 | 0.76 | 0.83 | 0.71 | 0.76 | 0.80 | 0.74 | 0.77 | 0.73 | 0.79 | 0.76 |
| GATK | C88 | Nonrepetitive | 0.78 | 0.75 | 0.76 | 0.61 | 0.60 | 0.60 | 0.50 | 0.49 | 0.49 | - | - | - | - | - | - |
| | | Repetitive | 0.66 | 0.54 | 0.59 | 0.48 | 0.42 | 0.45 | 0.41 | 0.37 | 0.39 | - | - | - | - | - | - |
| | Otava | Nonrepetitive | 0.76 | 0.72 | 0.74 | 0.57 | 0.47 | 0.52 | 0.48 | 0.18 | 0.26 | - | - | - | - | - | - |
| | | Repetitive | 0.65 | 0.57 | 0.61 | 0.45 | 0.31 | 0.36 | 0.40 | 0.11 | 0.17 | - | - | - | - | - | - |
| Varigraph | C88 | Bi-allelic | 0.80 | 0.81 | 0.80 | 0.72 | 0.79 | 0.75 | 0.74 | 0.78 | 0.76 | 0.65 | 0.72 | 0.69 | 0.76 | 0.83 | 0.80 |
| | | Multi-allelic | 0.88 | 0.73 | 0.80 | 0.74 | 0.66 | 0.70 | 0.71 | 0.69 | 0.70 | 0.78 | 0.71 | 0.74 | 0.82 | 0.80 | 0.81 |
| | Otava | Bi-allelic | 0.88 | 0.74 | 0.80 | 0.82 | 0.74 | 0.78 | 0.82 | 0.71 | 0.76 | 0.77 | 0.75 | 0.76 | 0.62 | 0.76 | 0.68 |
| | | Multi-allelic | 0.92 | 0.70 | 0.79 | 0.83 | 0.70 | 0.76 | 0.81 | 0.74 | 0.78 | 0.83 | 0.74 | 0.78 | 0.81 | 0.82 | 0.82 |
| GATK | C88 | Bi-allelic | 0.68 | 0.59 | 0.64 | 0.51 | 0.48 | 0.49 | 0.45 | 0.43 | 0.44 | - | - | - | - | - | - |
| | | Mult-iallelic | 0.50 | 0.40 | 0.45 | 0.28 | 0.23 | 0.25 | 0.16 | 0.18 | 0.17 | - | - | - | - | - | - |
| | Otava | Bi-allelic | 0.68 | 0.62 | 0.65 | 0.48 | 0.35 | 0.40 | 0.40 | 0.12 | 0.19 | - | - | - | - | - | - |
| | | Multi-allelic | 0.54 | 0.44 | 0.48 | 0.24 | 0.20 | 0.22 | 0.11 | 0.11 | 0.11 | - | - | - | - | - | - |

**Supplementary Table 17. Genotyping precision, recall and F-scores of Varigraph and GATK on autotetraploid potato genomes. Corresponding to Figure 5.**

See the file Supplementary.Table.17.xlsx
